# A novel injectable radiopaque hydrogel with potent properties for multicolor CT imaging in the context of brain and cartilage regenerative therapy

**DOI:** 10.1101/2023.04.20.537520

**Authors:** Moustoifa Said, Clément Tavakoli, Chloé Dumot, Karine Toupet, Yuxi Clara Dong, Nora Collomb, Céline Auxenfans, Anaïck Moisan, Bertrand Favier, Benoit Chovelon, Emmanuel Luc Barbier, Christian Jorgensen, David Peter Cormode, Danièle Noël, Emmanuel Brun, Hélène Elleaume, Marlène Wiart, Olivier Detante, Claire Rome, Rachel Auzély-Velty

## Abstract

Cell therapy is promising to treat many conditions, including neurological and osteoarticular diseases. Encapsulation of cells within hydrogels facilitates cell delivery and can improve therapeutic effects. However, much work remains to be done to align treatment strategies with specific diseases. The development of imaging tools that enable monitoring cells and hydrogel independently is key to achieving this goal. Our objective herein is to longitudinally study an iodine-labeled hydrogel, incorporating gold-labeled stem cells, by bicolor CT imaging after *in vivo* injection in rodent brains or knees. To this aim, an injectable self-healing hyaluronic acid (HA) hydrogel with long-persistent radiopacity was formed by the covalent grafting of a clinical contrast agent on HA. The labeling conditions were tuned to achieve sufficient X-ray signal and to maintain the mechanical and self-healing properties as well as injectability of the original HA scaffold. The efficient delivery of both cells and hydrogel at the targeted sites was demonstrated by synchrotron K-edge subtraction-CT. The iodine labeling enabled to monitor the hydrogel biodistribution *in vivo* up to 3 days post-administration, which represents a technological first in the field of molecular CT imaging agents. This tool may foster the translation of combined cell-hydrogel therapies into the clinics.

## 1. Introduction

Cell transplantation has become a promising approach to treat many debilitating conditions, in particular neurological and osteoarticular diseases.^[1]^ Various cell therapies have reached advanced clinical trial phases, many of which are based on mesenchymal stromal cells used to replace the lost cells and/or stimulate endogenous self-repairing mechanisms.^[2]^ However, one of the challenges still to overcome is the poor engraftment/survival of transplanted cells.^[3]^ This problem may be explained by untargeted delivery routes and low retention of the delivered cells *in situ*. Encapsulation of cells within hydrogels has been proposed as a promising strategy to circumvent these issues.^[4]^ Injectable hydrogels are attractive biomaterials that facilitate cell delivery and improve the effects of cell therapy due to their capability for homogeneous mixing with cells, their resemblance to biological tissues, and their minimally invasive administration.^[5]^ Beyond cell encapsulation, hydrogels themselves provide a favorable environment for tissue regeneration.^[4a]^ Therefore, the combination of hydrogel scaffold and stem cells may be referred to as a “repair kit”.

Hyaluronic acid hydrogels are appealing choices for cell encapsulation in a transplant approach. HA is a biocompatible and biodegradable polysaccharide ubiquitous in the body, and abundantly found in the brain and synovial fluid.^[6]^ Some work showed favorable effects of HA scaffolding on engrafted cell survival in preclinical models of stroke.^[7]^ However, the beneficial effect of cell-supporting HA hydrogel scaffolds is closely tied to the precision in injection, ability to fill irregular tissue defects and retention of the scaffold material within the tissue, which depend on the gelation properties of HA.^[8]^ Recently, we developed a new injectable HA hydrogel crosslinked by dynamic covalent bonds, suitable for cell delivery in a minimally invasive manner.^[9]^ Our hydrogel is produced by simply mixing two solutions of HA partners (HA modified with phenylboronic acid (PBA) and a fructose derivative (Fru)) in physiological conditions, allowing efficient and homogeneous cell encapsulation. This simple process of formation makes the hydrogel easily modifiable according to the targeted medical applications. Moreover, the dynamic covalent character of crosslinks (boronate ester bonds) in this hydrogel network makes it malleable, injectable and able to self-heal almost instantly.

Notwithstanding, impediments persist in the translation of hydrogel-based encapsulation technologies to clinical settings. Firstly, validation of the efficacy of repair kit delivery must be conducted on a per-subject basis, specifically to ascertain whether the lack of response to treatment is attributable to inadequate treatment potency or erroneous or substandard administration of treatment. Secondly, inquiry into the *in vivo* fate of hydrogel and cells is necessary to enhance the understanding of therapeutic mechanisms and to accordingly optimize treatment regimens. In pre-clinical assessments, hydrogels’ resemblance to the extracellular matrix poses challenges to their post-mortem identification on classical histological slices. Therefore, there is a crucial need for developing imaging tools that allow the distinct and simultaneous *in vivo* monitoring of cells on the one hand and cell-embedding hydrogels on the other hand.

We have recently demonstrated that the next generation CT, namely spectral photon-counting CT (SPCCT), was a method of choice for performing bicolor imaging of a dual-labeled repair kit in the context of translational stroke research, ^[10]^ where the hydrogel was made radiopaque by loading with iodinated nanoparticles. We have further shown that synchrotron K-edge subtraction CT (SKES-CT) provided a reference frame for SPCCT while providing higher spatial resolution and superior material differentiation.^[11]^ Interestingly, the biocompatibility and high stability in aqueous media of clinically-approved X-ray contrast agents make them compatible with hydrogels.^[12]^ Despite these advantages, the use of CT technology to longitudinally track hydrogels *in vivo* is limited by the technical challenges of fabricating biocompatible and biodegradable hydrogels exhibiting both long-persisting radiopacity and injectability. One of the limitations of our former study, for instance, was the fact that the hydrogel radiopacity was achieved via loading with iodinated nanoparticles, which may diffuse from the hydrogel, creating challenges for long-term follow-up.^[10]^ The covalent grafting of iodinated contrast moieties on the hydrogel-forming polymer is desirable to prevent the rapid leakage of contrast agents from the matrix.^[13]^ However, this procedure generally alters hydrogel properties, hence the necessity to find a trade-off between optimal X-ray opacity and physico-chemical properties of the HA scaffold.

In this context, this study aims: (i) to design and characterize a new iodine-labeled injectable HA (“HA-I”) hydrogel for therapeutic cell delivery with stable radiopacity and (ii) to demonstrate the ability to visualize the HA-I hydrogel *in vivo* with multimodal CT imaging in relevant rodent models in the context of brain and cartilage regeneration. For hydrogel labeling, our strategy was to functionalize the two HA partners with a clinical iodine-based contrast agent (i.e. 3-acetamido-2,4,6-triiodobenzoic acid, AcTIB). For *in vivo* testing, we chose to co-inject the HA-I hydrogel with human adipose-derived stromal cells (hASCs) labeled with gold nanoparticles (referred as the repair kit) and to image the dual-labeled “repair kit” thus obtained with bicolor SKES-CT (**Figure 1**).

**Figure 1.**
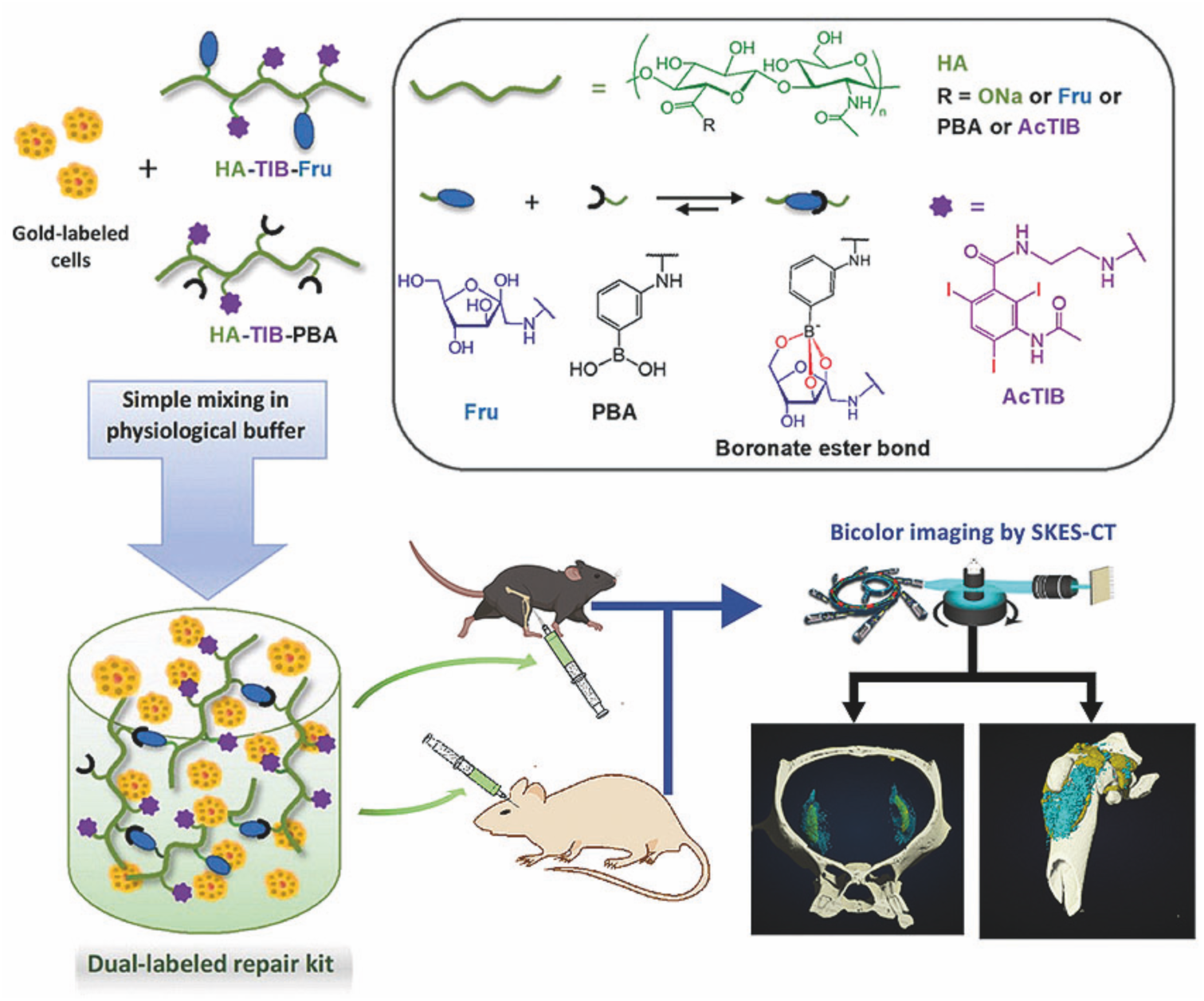
Schematic presentation of the strategy to design and synthesize the dual-labeled repair kit consisting of an iodine-labeled injectable HA hydrogel loaded with gold-labeled cells. The cell-laden hydrogel is produced by simply mixing two solutions of HA partners modified with the clinical iodine contrast agent AcTIB (HA-TIB-PBA and HA-TIB-Fru) in physiological conditions allowing efficient and homogeneous cell encapsulation. The dynamic covalent character of crosslinks (boronate ester bonds) in this hydrogel network makes it injectable and able to self-heal almost instantly. The *in vivo* examination of the HA-I alone and loaded with cells in knees of mice or brains of rats showed ability to longitudinally monitor by SKES-CT our dual-labeled repair kit.

## 2. Results

### 2.1. Synthesis and characterization of the iodine-labeled injectable HA hydrogel

The preparation of the iodine-labeled injectable HA hydrogel formulation required first the synthesis of the two HA hydrogel precursors, HA-TIB-Fru and HA-TIB-PBA, each labeled with a derivative of a clinical iodine-based contrast agent (AcTIB). Because of the very strong hydrophobicity of AcTIB moieties, the macromolecular parameters of the HA gel precursors (molar mass and degree of substitution (DS, average number of substituting groups per HA disaccharide unit)) were carefully chosen to ensure their perfect solubility in physiological conditions, and to obtain a hydrogel that shows appropriate rheological properties and easy injectability. The HA gel precursors were thus prepared from initial HA samples possessing different average molar masses (HA with M_w_ = 390 and 120 kg/mol, referred to as HA390 and HA120, respectively). HA390 and HA120 were first modified with *N*-(2-aminoethyl)-3-acetamido-2,4,6-triiodobenzamide (AcTIB-NH_2_) by an amide coupling reaction using 4-(4,6-dimethoxy-1,3,5-triazin-2-yl)-4-methylmorpholinium chloride (DMTMM) as a coupling agent and, DMTMM/HA and AcTIB-NH_2_/HA molar ratios of 0.36 and 0.60, respectively, to target DS of the HA conjugates of ∼ 0.2-0.3 (**Figure 2A**). AcTIB-NH_2_ was synthesized via amide linkage between *N*-Boc-ethylenediamine and AcTIB followed by removal of the *N*-Boc protecting groups (**Figure S1 and S2**, Supporting Information). Then, the HA390 and HA120 derivatives modified with TIB were reacted with 1-amino-1-deoxy-D-fructose (fructosamine) and 3-aminophenylboronic acid (APBA), respectively, using DMTMM for amide bond formation (**Figure 2B**). For the synthesis of HA-TIB-Fru, the DMTMM/HA and amine/HA molar ratios were fixed to 1 and 0.15, respectively, to obtain a DS_Fru_ of 0.15. Regarding that of HA-TIB-PBA, the amine/HA molar ratio was decreased to 0.1 to target a DS_PBA_ of 0.1 in order to maintain good water-solubility of the final HA conjugate.

**Figure 2.**
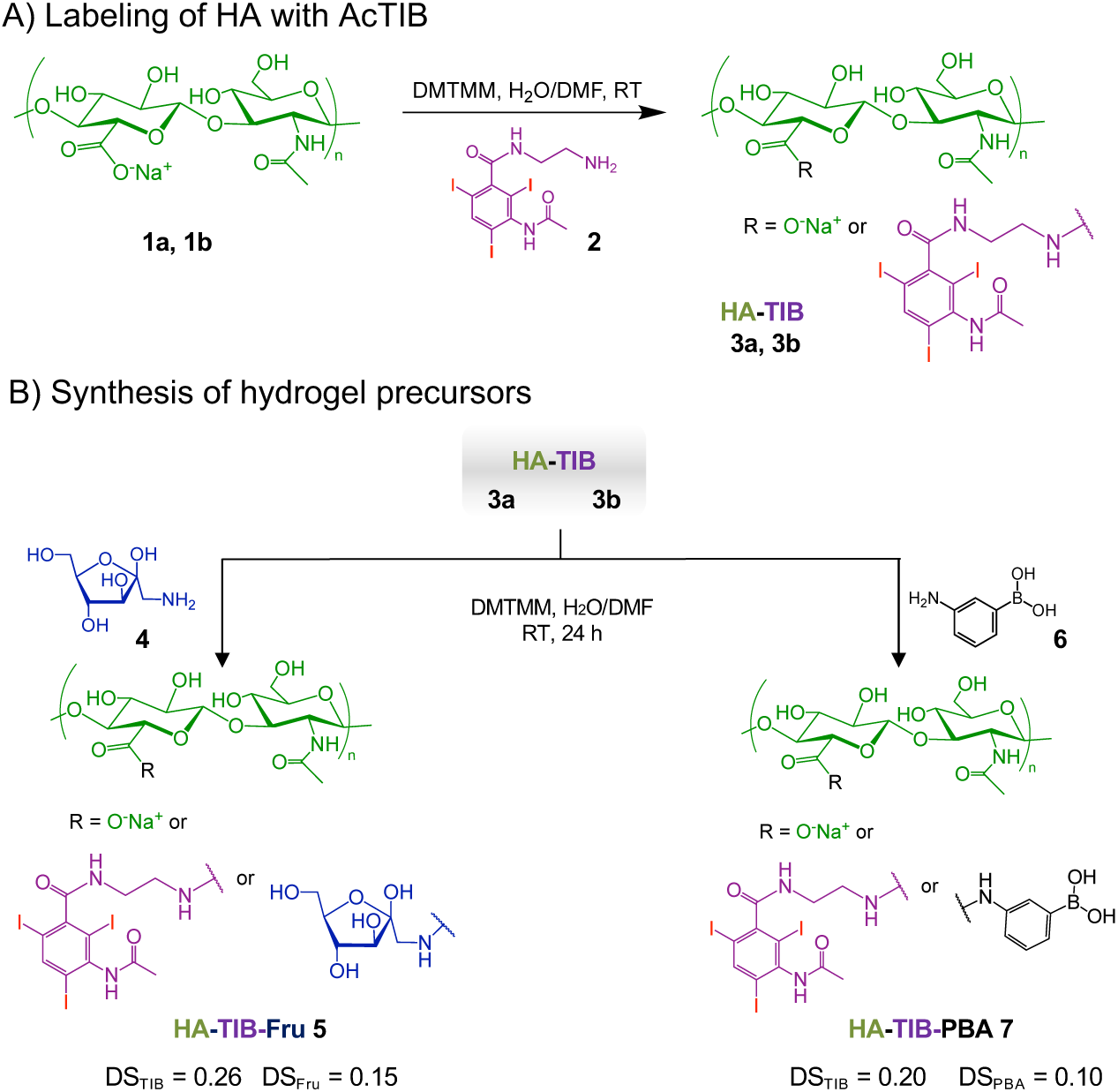
Synthesis of the iodine-labeled HA gel precursors. A) Modification of hyaluronic acid (HA390 **1a** and HA120 **1b** with an iodine-based contrast agent (AcTIB-NH_2_ **2**), affording HA-TIB **3a** and **3b**. B) Grafting of either fructosamine **4** or 3-aminophenylboronic acid **6** on HA-TIB to obtain HA-TIB-Fru **5** and HA-TIB-PBA **7**.

The chemical structures of the HA-TIB-Fru and HA-TIB-PBA derivatives were confirmed by ^1^H NMR spectroscopy (**Figure S3 and S4**, Supporting Information). Digital integration of the NMR spectra also allowed to assess their DS (DS_TIB_ = 0.25 and DS_Fru_ = 0.15 for HA-TIB-Fru, and DS_TIB_ = 0.20 and DS_PBA_ = 0.10 for HA-TIB-PBA).

Then, the iodine-labeled injectable HA (HA-I) hydrogel was produced by simply mixing thoroughly solutions of HA-TIB-Fru and HA-TIB-PBA in phosphate buffer saline (PBS, pH 7.4), at a total polymer concentration (*C_p_* = 18 g/L) and with a molar ratio of PBA-to-grafted fructose of 1. Dynamic rheological analyses revealed a gel-like behavior (G’ > G”) within the frequency window explored, as a result of formation of boronate ester cross-links between the two HA partners (**Figure 3A**). In addition, the storage moduli at 25 and 37 °C (G’_1Hz_ = 336 Pa and 249 Pa, respectively) were in the range of brain tissue stiffness (i.e., 200-2000 Pa)^[14]^. Because of their self-healing properties, boronate ester crosslinked HA hydrogels are excellent candidates for injectable hydrogel scaffolds that can be injected as preformed solid.^[9]^ In this regard, we evaluated the recovery of the HA-I hydrogel after large oscillatory strain deformations. As shown in **Figure 3B**, strain-dependent oscillatory measurements displayed a broad linear viscoelastic region with network failure at high strain (800 %). The network was shown to immediately recover its rheological properties when the strain was reduced to 10 %. Then, the gel was subjected to a series of two cycles of breaking and reforming, which consisted in applying large strain deformations (800 %), intercalated with low strain deformations (10 %) (**Figure 3C**). These strain-recovery experiments revealed full recovery of the gel network, demonstrating its self-healing property. Although dynamic rheological moduli and self-healing capacity are important parameters for determining injectability, injection tests of the HA-I hydrogel in an agarose brain phantom^[15]^ were also carried out to verify the suitability of the HA-I scaffold for intracerebral and intra-articular injection uses. To this end, the hydrogel (10 μL) was injected using a Hamilton syringe with a 26 G needle and a stereotaxic injector pump, similar to conditions used for intracerebral injections in rats. As illustrated in **Figure 3D** and in the video in Supporting Information (Video S1), the HA-I hydrogel stained in red (neutral red) could be injected with precision in the agarose brain phantom, at a rate of 5 µL/min.

**Figure 3.**
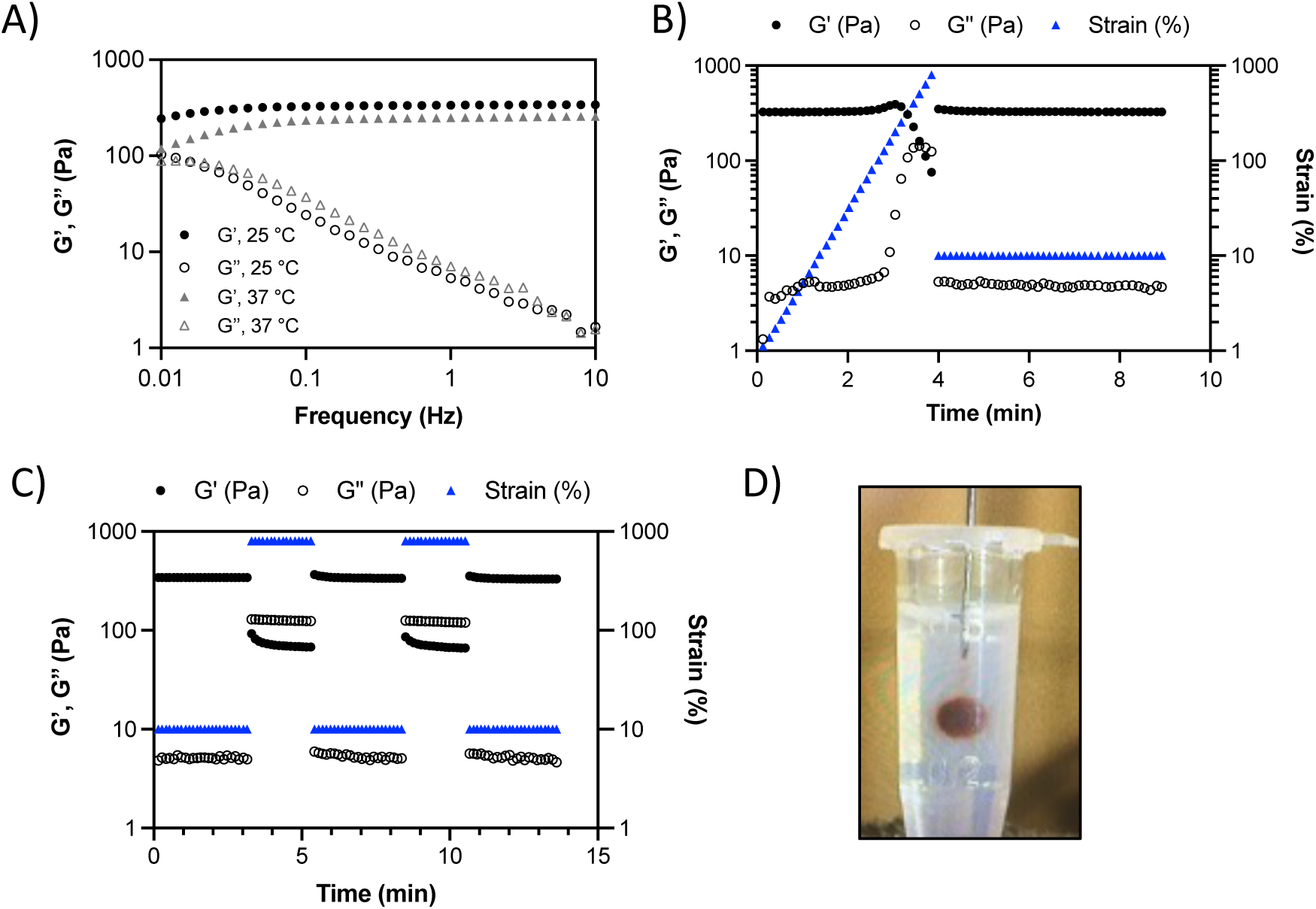
Rheological behavior and injectability of the HA-I hydrogel. A) Frequency dependence of the storage modulus (G’) and loss modulus (G’’) measured with 10 % strain at 25 °C and 37 °C. B) Variation of G’ and G’’ when increasing strain values to 800 % (hydrogel disruption), followed by reducing the strain to a constant value of 10 % (linear viscoelastic region). C) Alternate step strain sweep tests with alternating strain deformations of 10 and 800% at a fixed frequency (1 Hz). D) Photo of hydrogel injection in an agarose brain phantom through a 26 G (0.4 mm) needle (neutral red was added to color the hydrogel for visualization only).

### 2.2. Cell encapsulation and viability in the iodinated hydrogel

As a step towards injecting the HA-I hydrogel *in vivo*, we encapsulated hASCs within the hydrogel and investigated their viability for 3 and 7 days. In these experiments, the non-labeled HA hydrogel prepared by mixing HA-PBA and HA-Fru (referred to as “HA-ref hydrogel”, see experimental section and rheological behavior in Supporting information (**Figure S5**)) was used as a reference to assess the impact of the grafted AcTIB moieties on cell viability. Cells were encapsulated in the HA-I and HA-ref hydrogels at a density of 5×10^5^ cells/mL and cultured in growth media for 3 and 7 days (37 °C, 5 % CO_2_). As control, hASCs were incubated only in growth medium at a density of 5×10^3^ cells/mL (**Figure S6**, Supporting information). Thanks to the quasi-instantaneous gelation of the HA-TIB-Fru/HA-TIB-PBA mixture, the cells could be homogeneously distributed in 3D in the HA-I hydrogel, similar to the HA-ref hydrogel (**Figure 4A**). A live/dead assay was used to quantify the live (green) and dead (red) cells by fluorescence microscopy imaging of hASCs encapsulated in the iodine-labeled and non-labeled gel structure (**Figure 4B**).

**Figure 4.**
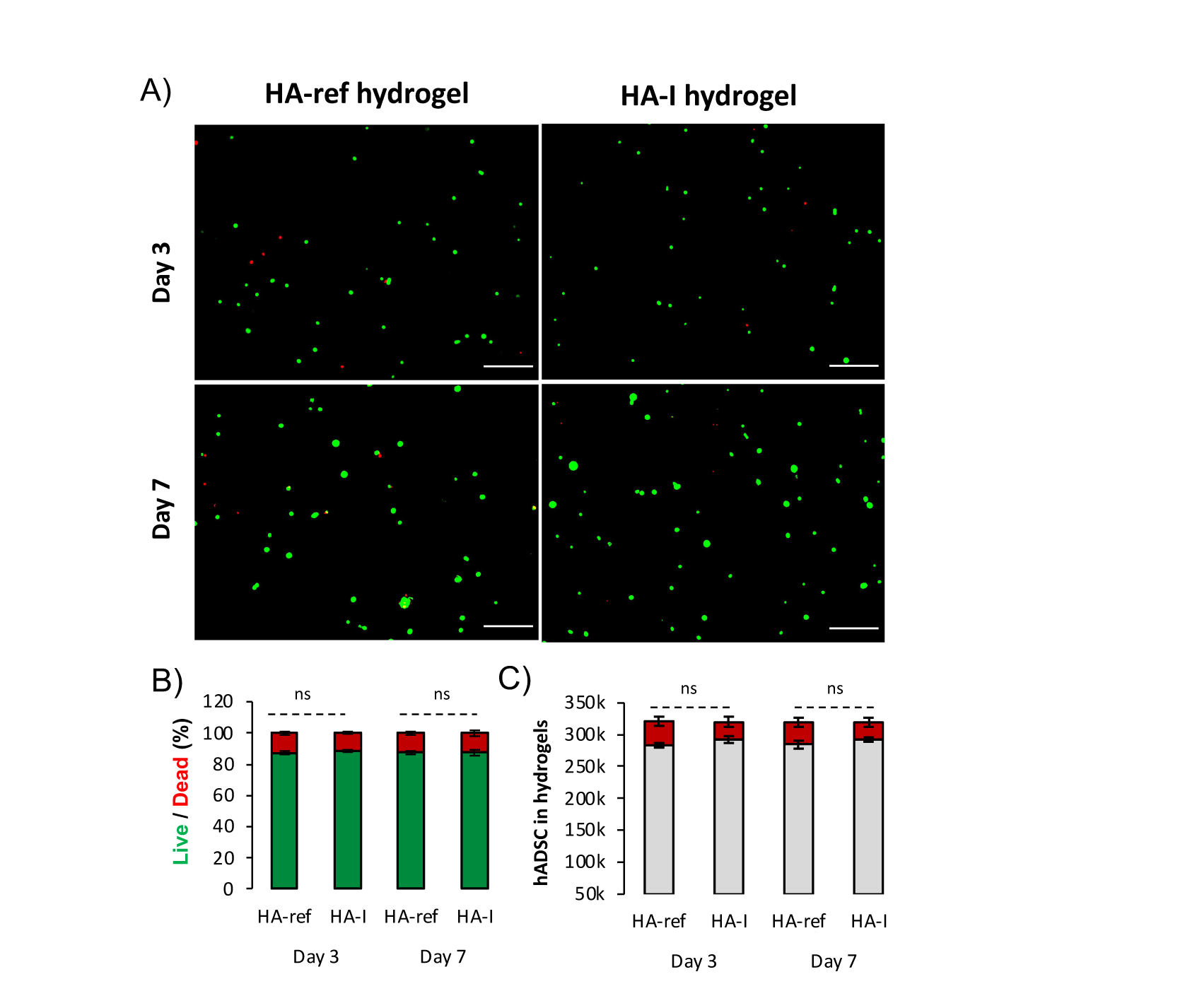
Cytocompatibility of the HA-I hydrogel in comparison to that of the HA-ref scaffold after 3 and 7 days of 3D cell culture. A) 2D microscopy images (fluorescence, scale bar = 200 μm) of Live/Dead (green/red) staining of hASCs encapsulated in the HA-ref and HA-I hydrogels after 3 and 7 days of culture. Quantification of cell viability from B) Live/Dead assay and C) trypan blue assay, represented as mean ± SD (*n* = 3). Statistical analysis from one-way ANOVA test, * indicating p< 0.05; ns: not significant.

The HA-I hydrogel showed high cytocompatibility over the 7 days of cell culture (> 80 % of cell viability), similar to the HA-ref hydrogel. This result was confirmed using the trypan blue dye exclusion assay (**Figure 4C**). These findings reveal not only the ability of the HA-I hydrogel to encapsulate cells in 3D, but also to provide a microenvironment conducive to maintain cellular survival over one week.

### 2.3. *In vitro* SKES-CT imaging of HA-I hydrogel

Next, we examined the ability to visualize the HA-I hydrogel by bicolor imaging with SKEST-CT *in vitro*. To this end, hydrogels with embedded cells were prepared by mixing both iodine labeled HA gel precursors (HA-TIB-Fru and HA-TIB-Fru) resulting in an iodine concentration of 2.4 mg/mL ([HA-I hydrogel] = 18 g/L) and gold-labeled hASCs at different concentrations. Four tubes containing the dual-labeled repair kit consisting of 10 µL of HA-I hydrogel loaded with a range of AuNPs-labeled hASCs (1.25×10^5^, 2.5×10^5^, 3.75×10^5^, 5×10^5^) were prepared. Tubes were imaged with SKES-CT and compared to a control tube that contained only agarose gel (**Figure 5**). On conventional attenuation images, moderate enhancement was seen from HA-I hydrogel compared to agarose (**Figure 5A**). However, the iodine maps clearly depicted HA-I hydrogel in all 4 tubes (**Figure 5B**). AuNP-labeled hASCs were also clearly visible on the gold maps, even for the smallest quantity of 1.25×10^5^ cells (**Figure 5C-D**).

**Figure 5.**
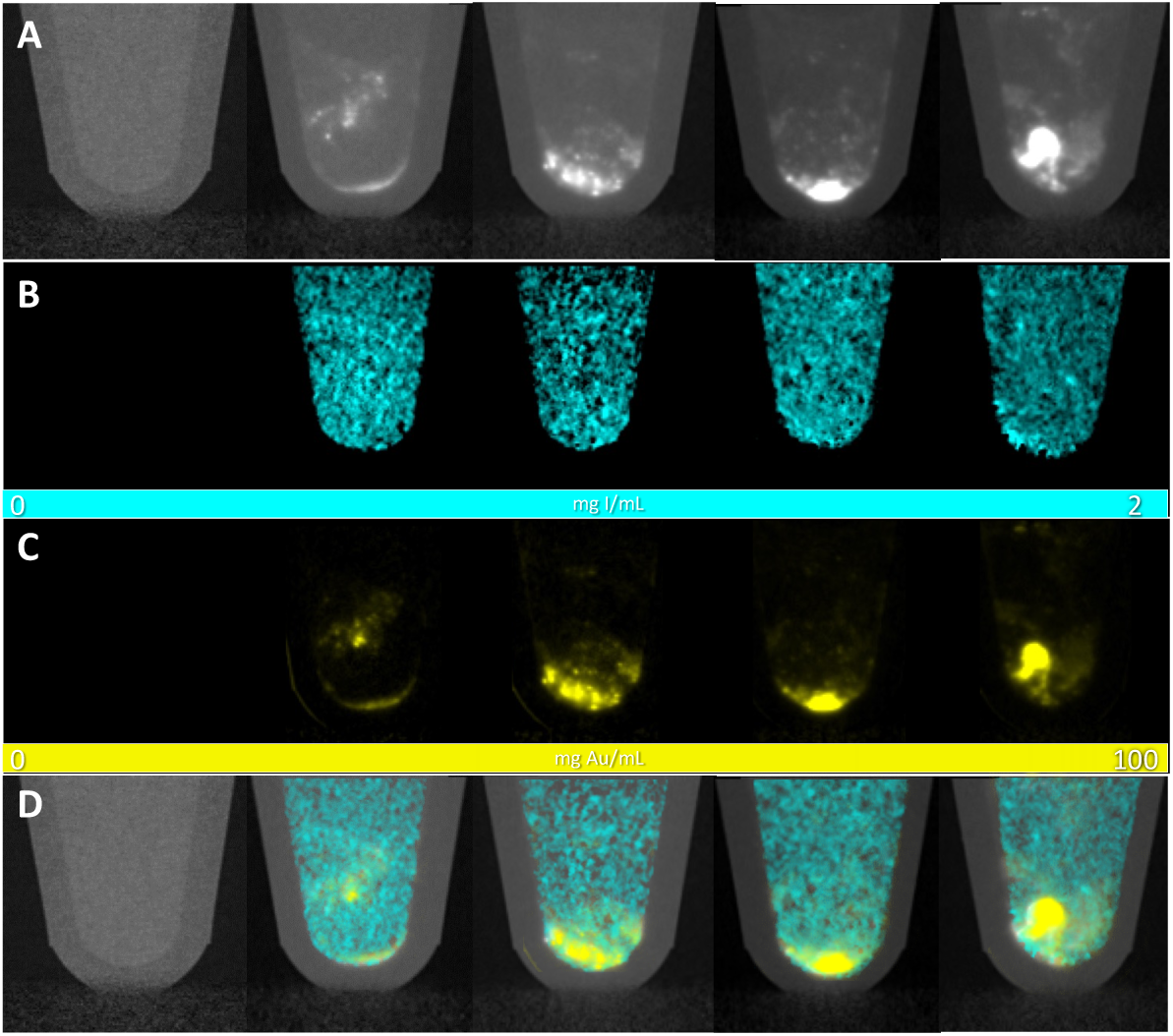
*In vitro* imaging of dual-labeled repair kits with SKES-CT. A) Attenuation images of tubes containing gold-labeled hASCs encapsulated in HA-I hydrogel (representative single slice from 3D data set). Iodine and gold (and therefore hydrogel and hASCs) cannot be distinguished on these conventional images. B) Iodine concentration maps highlighting hydrogel distribution. C) Gold concentration maps of the tubes highlighting AuNPs-labeled hASCs distribution. D) Attenuation images superimposed with iodine and gold concentration maps. From left to right: Agarose tube, tubes containing 1.25×10^5^, 2.5×10^5^, 3.75×10^5^ and 5×10^5^ AuNPs-labeled hASCs encapsulated in HA-I hydrogel.

### 2.4. *In vivo* studies

Given the radiopaque properties of the HA-I hydrogel, the next step consisted in imaging the repair kit *in vivo* after *in-situ* administration in 2 organs of interest with regard to cell therapy: the brain and the knee. Experiments at the synchrotron were organized in 2 sessions: in the first one, brain and knee samples were imaged *ex vivo* to ascertain the feasibility of the imaging approach before involving live animals (in line with 3R principles) and the second session was dedicated to *in vivo* imaging of a mouse model of osteoarthritis.

#### 2.4.1. Imaging of the HA-I hydrogel administered in healthy rat brains

Our first aim was to evaluate the potential of the HA-hydrogel for cell delivery and imaging for neurological applications. The dual labeled repair kit consisted in 10 µL HA-I hydrogel loaded with 2.5×10^5^ AuNP-labeled hASCs. Four healthy rats were intracerebrally injected with different combinations of the repair kit components as detailed in Table 1.

**Table 1.** Administration pattern for the four rats that received an intracerebral administration of the repair kit and/or its individual components. ID: identification. The dual-labeled repair kit consisted in 10-µL HA-I hydrogel loaded with 2.5×10^5^ AuNP-labeled hASCs.

| Rat ID | Left hemisphere | Right hemisphere |
| --- | --- | --- |
| 1 | Dual-labeled repair kit | $2.5 \times 10^5$ AuNP-labeled hASCs |
| 2 | Dual-labeled repair kit | HA-I hydrogel |
| 3 | $2.5 \times 10^5$ AuNP-labeled hASCs | HA-I hydrogel |
| 4 | Dual-labeled repair kit | Dual-labeled repair kit |

##### 2.4.1.1. Preclinical micro-CT imaging

We first assessed whether we could monitor the repair kit delivery using conventional micro-CT (micro-CT), immediately after administration. Although only slight enhancement was seen on attenuation images, HA-I hydrogel delivery could be readily visualized *via* micro-CT using the appropriate window (**Figure 6A** and **6C**, blue arrow: HA-I hydrogel alone and green arrow: HA-I hydrogel in dual-labeled repair kit). However, these conventional images did not allow to discriminate the HA-I hydrogel and the AuNP-labeled hASCs. For this reason, we only segmented the HA-I hydrogel when injected alone (**Figure 6B** and **6D**). This segmentation gave a mean hydrogel volume of 9.8 ± 0.5 μL (N=2), in excellent agreement with the 10 µL volume injected.

**Figure 6.**
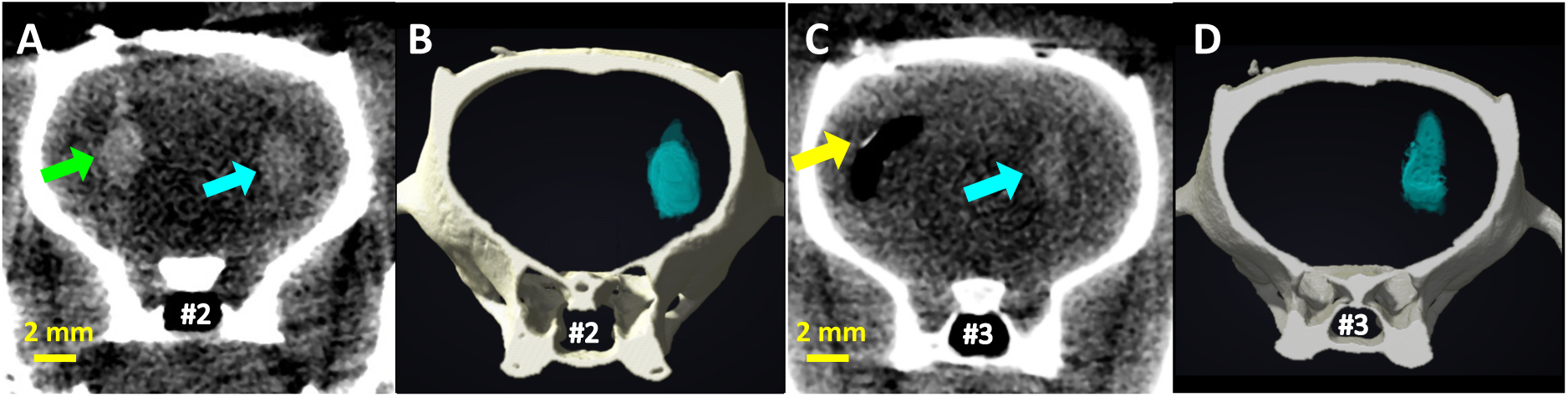
Imaging of the dual-labeled repair kit and its individual components with micro-CT in two healthy rat brains (rats #2 and #3). A) and C) Attenuation images of each brain (representative single slice from 3D data set). B) and D) 3D view of the segmented bone (white) and HA-I hydrogel administered alone (blue) for volume quantification purpose. Green arrow: repair kit, blue arrows: HA-I hydrogel alone, and yellow arrow: AuNPs-labeled hASCs alone.

##### 2.4.1.2. *Ex vivo* SKES-CT imaging

To investigate the ability to visualize the HA-I hydrogel by bicolor imaging in a rat model, the same brains were then imaged with SKES-CT after their sacrifice at 72 h post-injection (**Figure 7**). Interestingly, iodine in the HA hydrogel was still present in the conventional images (**Figure 7**, blue and green arrows), indicating stability of labeling. As expected, iodine enhancement was better seen in the conventional images of SKES-CT compared to that of micro-CT (**Figure 7A**). Yet, as for micro-CT, these conventional images did not allow the differentiation of the HA-I hydrogel from the AuNP-labeled hASCs. In contrast, bicolor imaging allowed to discriminate the HA-I hydrogel (**Figure 7B**) from the AuNPs-labeled hASCs (**Figure 7C**). The distribution of AuNPs-labeled hASCs was more homogenous within the hydrogel when administered *in vivo* than *in vitro* (**Figure 7D**), since the high volume of hydrogel used (1 mL) for preparing the repair kit for preclinical studies makes the homogeneous mixing of hASCs and hydrogel easier. The mean volume occupied by the HA-I scaffold calculated using the segmented 3D volumes (**Figure 7D**) was 10.2 ± 1.5 μL (N=6). This value was in excellent agreement with the actual volume of injection (10 µL). This suggests that the hydrogel had not been eliminated nor degraded at 3 days post-administration in the healthy brain.

**Figure 7.**
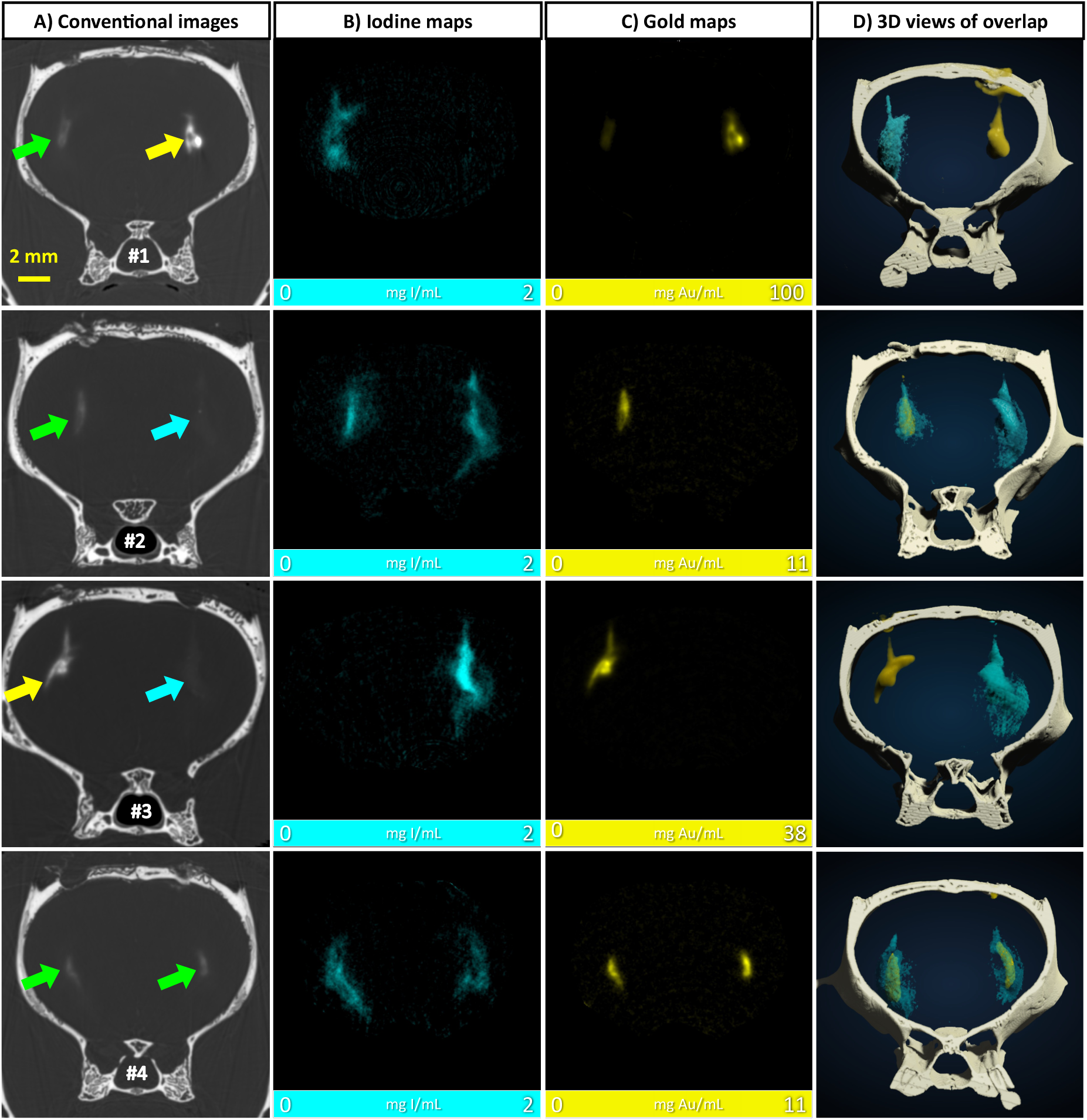
Imaging of the dual-labeled repair kit and its individual components with SKES-CT in the brain of four healthy rats. Results for each brain are displayed on each row. A) Attenuation images (representative single slice from 3D data set). Green arrows: repair kit, yellow arrows: AuNPs-labeled hASCs alone and blue arrows: HA-I hydrogel alone. B) Corresponding iodine concentration maps. C) Corresponding gold concentration maps. D) 3D view of segmented bone (white), iodine (blue) and gold (yellow).

#### 2.4.2. Imaging of the HA-I hydrogel administered in the knees of healthy and osteoarthritic mice

##### 2.4.2.1. *In vivo* SKES-CT imaging

In order to further explore the potential of the HA-I hydrogel for both cell-delivery and imaging in the treatment of chronic diseases, we investigated its behavior after intra-articular injection in mouse knees. In a first experiment, the dual-labeled repair kit was injected in the healthy knee joints of three mice. The dual labeled repair kit consisted of 2.5 µL HA-I hydrogel loaded with 2.5×10^5^ (N=2) or 3.5×10^5^ (N=1) AuNP-labeled hASCs. Mice were sacrificed immediately after administration and bi-color imaging with SKES-CT was performed *ex vivo*. Images showed the presence of gold and iodine inside knee joints in all mice (**Figure 8**), demonstrating that the HA-I hydrogel can be used to monitor intra-articular delivery to the target site. AuNP-labeled hASCs appeared homogenously distributed within the hydrogel (**Figure 8D**). The mean volume occupied by the HA-I scaffold calculated using the reconstructed 3D images (**Figure 8D**) was 2.7 ± 1.4 μL (N=3). This value was in excellent agreement with the actual volume of injection (2.5 µL).+

**Figure 8.**
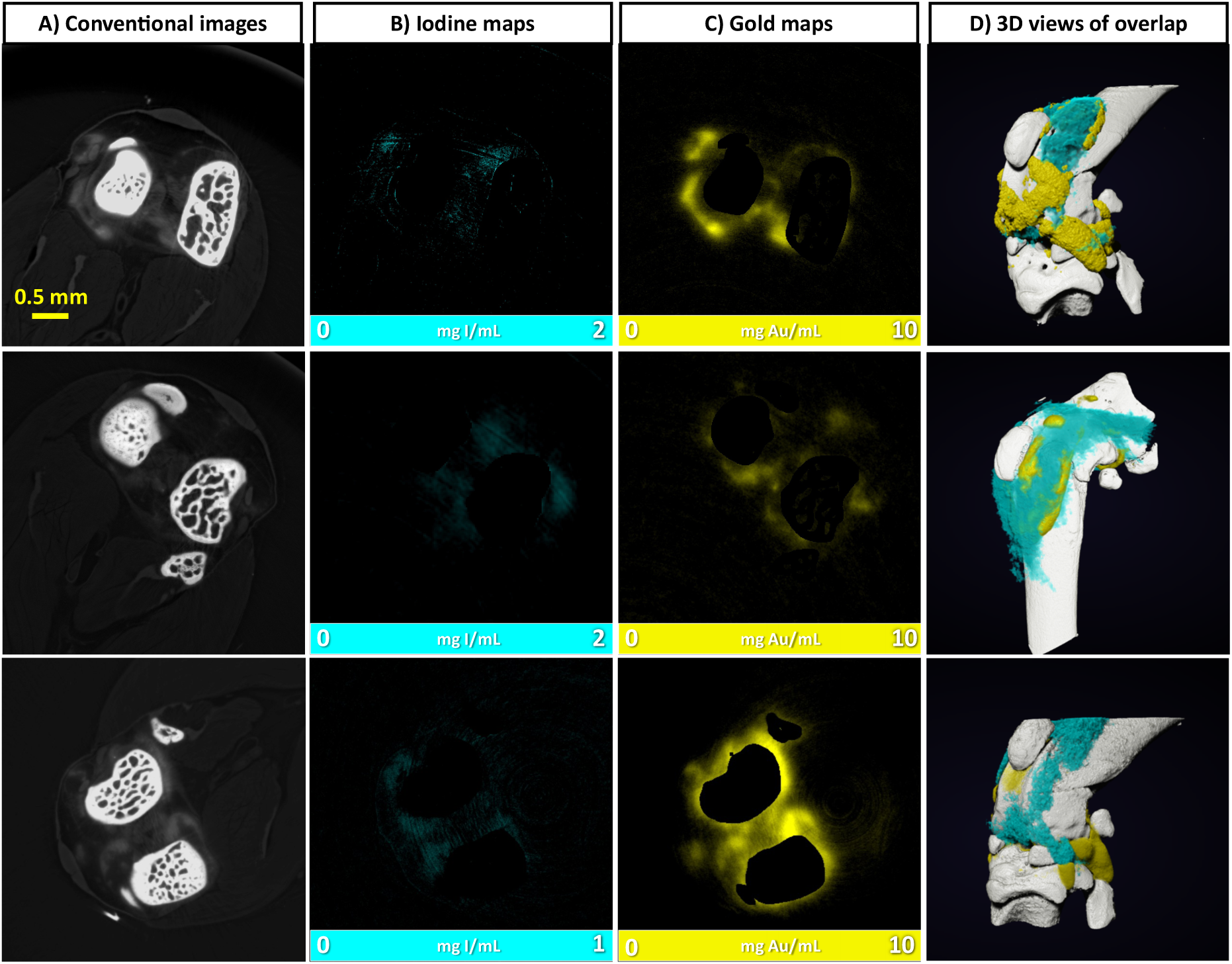
Imaging of the dual-labeled repair kit with SKES-CT in the knees of healthy mice. Results for each knee are displayed on each row. A) Attenuation images (representative single slice from 3D data set). B) Corresponding iodine concentration maps. C) Corresponding gold concentration maps. D) 3D view of segmented bone (white), iodine (blue) and gold (yellow).

In the next experiment, we aimed to evaluate our imaging approach in a mouse model of osteoarthritis. The HA-I hydrogel was injected in the knees of osteoarthritic mice (N=11) and its distribution was investigated in live mice in the first three days post-administration (**Figure 9**, 24h: N=2, 48h: N=4 and 72h: N=5). The dual-labeled repair kit consisted in 2.5 µL HA-I hydrogel loaded with 2.5×10^5^ AuNP-labeled hASCs. The iodine signal was present in 100% of knees at 24h (2/2), 75% at 48h (3/4) and 60% at 72h (3/5). Considering the stability of iodine labeling as seen in brain experiment, this suggests that the scaffold was eliminated from the joint in the very first days following its administration. The mean volume occupied by the HA-I scaffold *in vivo*, calculated using the reconstructed 3D images (**Figure 9**) was 1.8 ± 1.0 μL (N=11). This value was in line with the actual volume of injection (2.5 µL) considering the fact that in this case, some elimination had already occurred.

**Figure 9.**
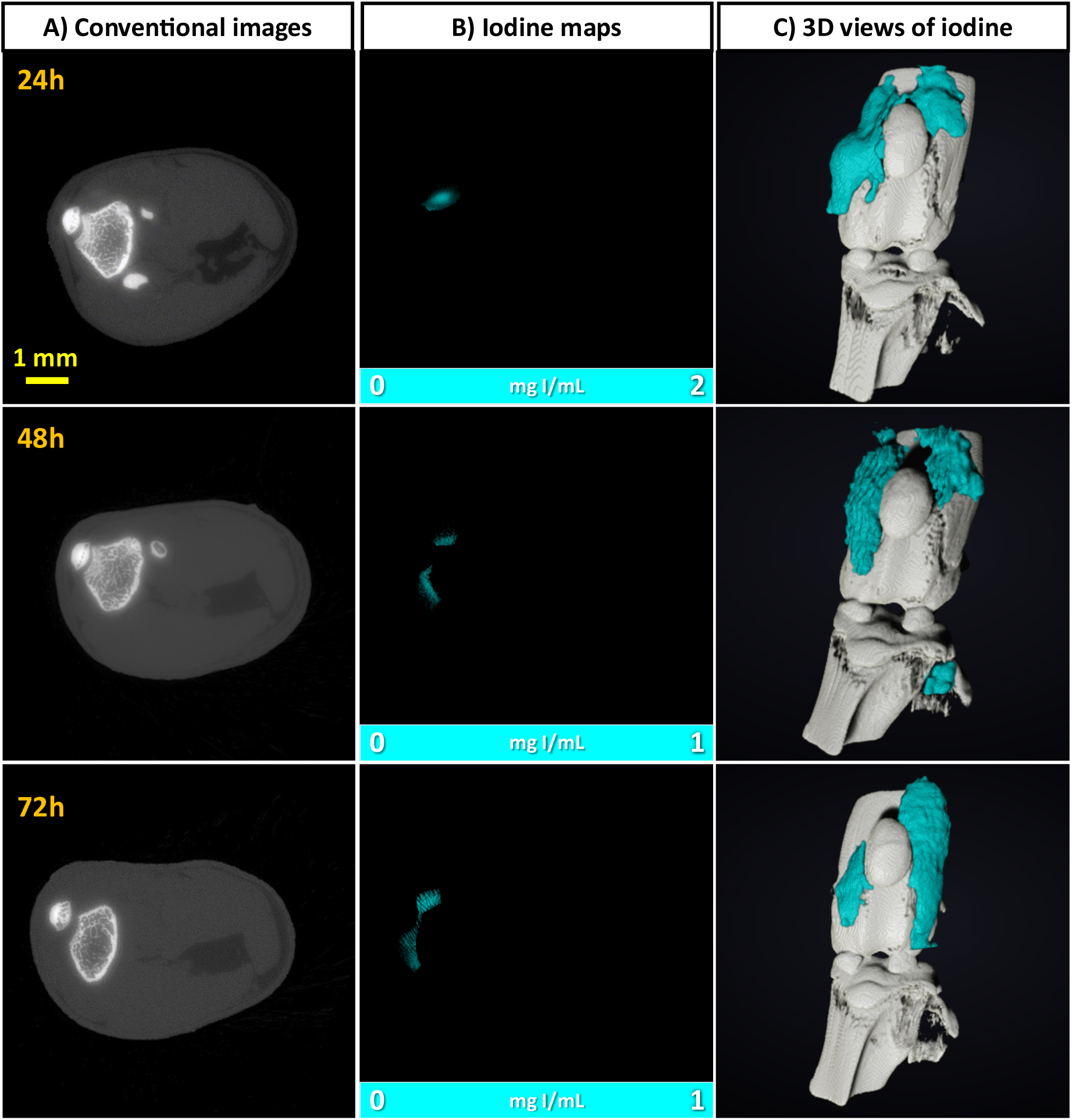
Imaging of the HA-I hydrogel with SKES-CT in the knees of osteoarthritic mice. Results of 3 representative knees imaged at 3 different times post-administration are displayed on each row. A) Attenuation images (representative single slice from 3D data set). B) Corresponding iodine concentration maps. C) 3D view of segmented bone (white) and iodine (blue).

##### 2.4.2.2. *Ex vivo* X-ray phase contrast tomography (XPCT)

Last, we sampled the knees of mice and imaged them post-mortem with synchrotron X-ray phase contrast tomography, in order to obtain a 3D Phase Contrast image of the knee joints taken as ground truth at the spatial resolution of 6 µm. The hydrogel distribution could be thus readily depicted within the joint (**Figure 10**, Video S2) and was in agreement with SKES-CT findings. This further validated iodine labeling stability in time and detectability with SKES-CT.

**Figure 10.**
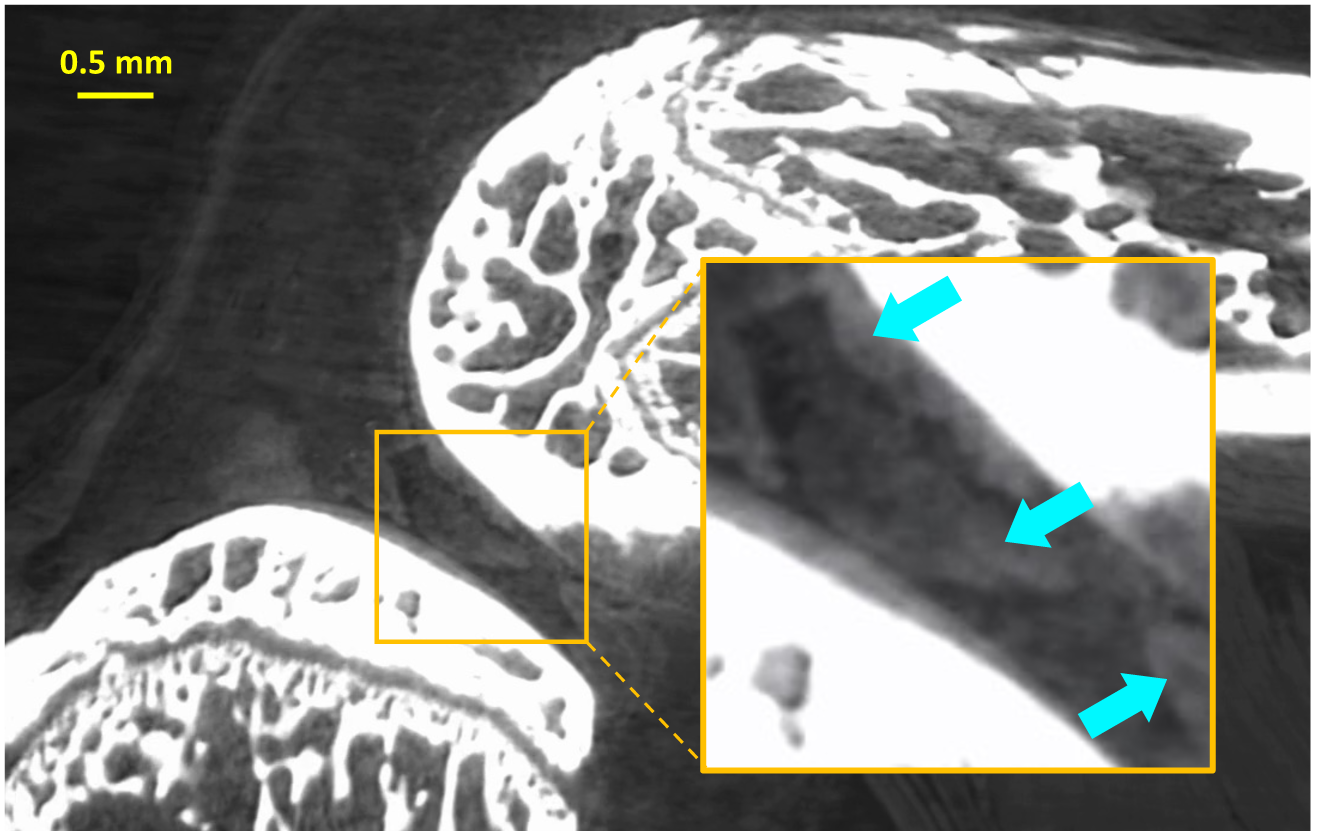
Imaging of the injected HA-I hydrogel with XPCT in the osteoarthritic mouse knee. The hydrogel can be identified in the joint capsule thanks to the iodine labeling (indicated by the blue arrows and Video S2).

## 3. Discussion

In this study, we designed and characterized a novel iodine-labeled injectable self-healing HA hydrogel for stem cell delivery. We first demonstrated that iodine labeling did not alter the rheological, self-healing and injectability properties of the hydrogel. We also verified *in vitro* that iodine labeling did not impact 3D encapsulation and viability of hASCs. Next, we aimed to establish that X-ray radiopacity provided by iodine labeling was sufficient for imaging the gel with CT. Bicolor CT imaging is emerging as a promising technique for tracking cells and hydrogel simultaneously.^[10]^ We therefore labeled hASCs with gold nanoparticles, encapsulated them within the iodine-labeled HA hydrogel, and imaged the formed dual-labeled repair kit with bicolor SKES-CT *in vitro* and *in vivo*. Our study provides the proof-of-concept that iodine labeling enabled to monitor the hydrogel biodistribution *in vivo* in rodent brains and knees up to 3 days post-administration of the dual-labeled repair kit. These organs were chosen because neurological and osteoarticular diseases are considered prime applications for stem cells-based therapies. The volume of hydrogel could be estimated thanks to the 3D reconstructions of CT scans. Interestingly, the *in vivo* fate of gel and cells differed according to the target tissue. Taken together, our data indicate that the iodine-labeled HA hydrogel that we have designed presents stable iodine labeling as well as excellent properties for cell encapsulation and injection. In addition, our gel is suitable for imaging with translational CT, from bench-top micro-CT to cutting-edge spectral CT. To the best of our knowledge, this represents a technological first in the field of molecular CT imaging agents.

Hydrogels are the most widely used scaffolds in regenerative medicine.^[16]^ Beside biocompatibility and degradability, injectability is the most important characteristic required for the hydrogel scaffold to be delivered through minimally invasive procedures. To be injectable, the hydrogel can be either a pre-crosslinked network that undergoes a shear-thinning process during injection or a liquid formulation that is gelled *in situ* using environmental triggers (pH, temperature, ionic strength), a chemical crosslinker or a click coupling reaction.^[17]^ However, there are some potential drawbacks to the *in-situ* crosslinking approaches, which include potential adverse effects on cell viability and/or low gelation kinetics. Rapid gelation of the pre-polymer solution is indeed a prerequisite for achieving cell retention at the target site. Injectable self-healing hydrogels, assembled through dynamic covalent bonds, are a promising solution for hydrogel-assisted cell delivery as they afford the unique property of being injectable even after having formed a gel. Indeed, their solid-like properties within the syringe allows to maintain cells homogeneously suspended, leading to more reproducible and consistent cell delivery.^[18]^ In the current study, the dynamic covalent nature of the crosslinking bonds (boronate ester bonds) between HA chains, makes the hydrogel suitable as an injectable scaffold with rapid self-healing capability.

Using hydrogel as a combined drug-delivery and imaging (theranostic) platform is a growing field of interest.^[19]^ There are been some attempts in the literature to label hydrogels by simply mixing them with iodinated contrast agent or nanoparticles; however, this resulted in loss of the labeling agent in the first days following administration.^[10, 13b]^ For hydrogel labeling, our strategy was to functionalize the two HA gel precursors with a clinical iodine-based contrast agent (AcTIB) through covalent bonds for long-term visualization. Regarding clinical iodine contrast agents, those used today are all based on tri-iodinated benzene rings that tend to increase the hydrophobicity of polymers. This makes hydrogel labeling via covalent linkage of iodine moieties technically challenging, as this can change the polymer network properties. This difficulty was underlined by Lei et al. who developed an *in-situ* gelling polymer system with long-acting radiopacity for the first time.^[20]^ This hydrogel consists of a thermogelling system based on a 2,3,5-triiodobenzoic acid (TIB)-capped diblock copolymer of methoxy poly(ethylene glycol) and poly(lactic acid). The covalent introduction of TIB moieties enabled *in vivo* monitoring of the hydrogel administered in the abdomen to prevent post-operative abdominal adhesions with micro-CT in a rat model.^[20a]^ Yet, to the best of our knowledge, no injectable dynamic covalent hydrogel scaffolds with good and stable radiopacity has been reported so far. In the present study, such a hydrogel, entirely made of HA, could be successfully obtained by carefully choosing the concentration of iodine contrast agent in the polysaccharide network. We found iodine labeling conditions that did not alter the biocompatibility of the HA hydrogel, neither its rheological properties (G’ ∼ 336 Pa). This G’ modulus is suitable for brain repair because it matches brain mechanical properties.^[7a, 21]^ It is also in the range of the G’ values of HA hydrogels used for knee viscosupplementation.^[22]^ Viscosupplementation is a clinically-approved treatment of osteoarthritis that consists in injecting a HA gel into the knee joint. Adding HA to the arthritic joint (without stem cells) is expected to facilitate movement and reduce pain. However, some patients are not relieved by the injections. Therefore, the iodine-labeled HA hydrogel that we here introduce may also help understand the outcome of viscosupplementation treatment in osteoarthritis.

Importantly, the AcTIB moieties attached to the HA hydrogel did not induce cytotoxic effect to hASCs, as shown by the limited amount of dead cells after 7 days of cell culture, similar to the non-labeled HA hydrogel. hASCs were used because they are an abundant and accessible source of adult stem/stromal cells and hold no ethical concerns. Moreover, the efficacy of hASCs has been confirmed in the pre-clinical treatment of many disorders and injuries, including ischemic stroke,^[23]^ neurodegenerative disorders,^[24]^ cartilage damage,^[25]^ rheumatoid arthritis,^[26]^ and osteoarthritis^[27]^. In addition, hASCs have already been evaluated in Phase I and II clinical trials for ischemic stroke^[28]^ and knee osteoarthritis^[29]^.

The radiopaque HA-I hydrogel could be readily visualized by micro-CT imaging despite the fact that this modality is not very sensitive. While this conventional CT method can be used to report on the delivery of the hydrogel, it cannot be applied for the simultaneous and selective visualization of the hydrogel and cells. This could be achieved only by using bi-color imaging with SKES-CT. In the current study, we demonstrated that iodine-labeling of hydrogel was valuable for the tracking of the repair kit with this innovative imaging modality. This imaging approach based on dual labeling has the potential to provide a more effective and efficient evaluation of outcomes of hydrogel-assisted cell therapies by relating treatment efficacy to the fate of therapeutic cells together with their carrier. As SKES-CT has the capacity to reconstruct three dimensional (3D) volume of implanted materials deep within the body and is intrinsically quantitative,^[30]^ it can provide accurate information about their *in vivo* behavior and long-term degradation. Post-mortem high resolution synchrotron X-ray phase contrast tomography confirmed the 3D distribution of iodine-labeled HA gel in the joint knee.

In this regard, the 3D reconstruction images provided evidence that the HA-I hydrogel was precisely injected at the target site in the brains of rats and in the knee joints of mice. In addition, they showed that the volume of the HA-I gel remained unchanged at the injection site in the first 3 days following administration in the healthy brain. However, hydrogel partial elimination was observed in the same time-lapse following administration in a mouse model of osteoarthritis. The difference might be due to two factors. First, the fact that mice were physically active after administration: the mechanical constraints induced by joint mobilization may have favored hydrogel elimination, while in turn, in the brain, the absence of movement may favor long-term remanence. Second, the fact that the osteoarthritic knee is a highly inflammatory environment. Macrophages may actively participate in hydrogel elimination in this case, while in the healthy knee and brain, the hydrogel might remain on the long-term due to the absence of inflammatory condition. These hypotheses remain to be tested in further studies specifically designed to address these issues (healthy vs damaged brains/knee joints with a long-term follow-up).

## 4. Conclusion

In the present study, we developed a radiopaque self-healing hydrogel exclusively composed of HA through a simple procedure allowing homogeneous encapsulation of hASCs. Its solid-like properties within the syringe allowed to maintain cells homogeneously suspended. Additionally, its autonomous self-healing characteristics enabled their accurate delivery at the targeted site as observed for the first time by bi-color imaging with SKES-CT. The volume of the hydrogel calculated on the basis of 3D reconstruction provided valuable information about precision of injection and stability/elimination of the gel 3 days post-administration taking advantage of its long-acting radiopacity. All together, these results demonstrate that the combination of cell labeling and iodine-labeled hydrogel with SKES-CT can foster the translation of cell therapy for the benefit of patients with osteoarthritis and ischemic stroke, which are leading cause of disabilities. Future studies will focus on translating this theranostic approach using SPCCT for translational purposes, first with a pre-clinical SPCCT. Simultaneous monitoring of cells and gels open an avenue for better understanding stem cells-based therapies and promoting their clinical translation.

## 5. Experimental Section

### Materials

Hyaluronic acid sodium salt samples possessing a weight-average molar mass (M_w_) of 390 and 120 kg/mol were purchased from Contipro France. The molar mass distribution and the weight-average molar mass of these samples were determined by size exclusion chromatography using a Waters GPC Alliance chromatograph (USA) equipped with a differential refractometer and a light scattering detector (MALS) from Wyatt (USA); the solution was injected at a concentration of 1 mg/mL in 0.1 M NaNO_3_, at a flow rate of 0.5 mL/min and at a column temperature of 30 °C. The dispersity (*Đ*) of the samples is M_w_/M_n_ ≈ 1.5-2. 1-Amino-1-deoxy-D-fructose hydrochloride (fructosamine) was supplied by Carbosynth. 3-Aminophenylboronic acid hemisulfate salt (APBA), 4-(4,6-dimethoxy-1,3,5-triazin-2-yl)-4-methylmorpholinium chloride (DMTMM), phosphate-buffered saline (PBS), 3-acetamido-2,4,6-triodobenzoic acid bis(2-hydroxyethyl)-ammonium salt, *N*-Boc-ethylenediamine, 1-[bis(dimethylamino)methylene]-1H-1,2,3-triazolo(4,5-b)pyridinium 3-oxide hexafluorophosphate (HATU), 11-mercaptoundecanoic acid, gold (III) chloride hydrate, sodium citrate dihydrate, agarose (Reference A9539) other chemicals were purchased from Sigma-Aldrich and were used without further purification. For in vitro cell cytocompatibility studies and the gold labeling procedure, therapeutic grade human mesenchymal adipose-derived stem cells were provided from EFS (“Etablissement Français du Sang”) for *in vitro* experiments, “Banque de tissus et cellules Hospices Civils de Lyon” for brain experiments and ECell France for knee experiments. Live/Dead kit staining were purchased from Merk. Lysat plaquet and heparin 5000 U/mL, beta fibroblast growth factor (βFGF), 3-(4,5-dimethylthiazol-2-yl)-2,5-diphenyl tetrazolium bromide (MTT), Dulbecco’s phosphate buffer saline, ⍺-MEM (⍺-Minimum Essential Media), Trypan Blue were purchased from ThermoFisher Life Science. *N*-(2-aminoethyl)-3-acetamido-2,4,6-triiodobenzamide (AcTIB-NH_2_, **2**) was synthesized as described in Supporting Information (**Figure S1**). Non-labeled HA-PBA (DS_PBA_ = 0.15) and HA-Fru (DS_Fru_ = 0.15) were prepared from HA with M_w_ = 390 kg/mol as described previously^[9]^.

### Methods

#### NMR spectroscopy

^1^H NMR spectra were recorded at 25 °C or 80 °C using a Bruker AVANCE III HD spectrometer operating at 400.13 MHz (^1^H). ^1^H NMR spectra were recorded by applying a 90° tip angle for the excitation pulse, and a 10 s recycle delay for accurate integration of the proton signals. Deuterium oxide (D_2_O) and deuterated dimethylsulfoxide (DMSO-d6) were obtained from Euriso-top (Saint-Aubin, France). Chemical shifts (δ in ppm) are given relative to external tetramethylsilane (TMS = 0 ppm) and calibration was performed using the signal of the residual protons of the solvent as a secondary reference. All NMR spectra were analyzed with Topspin 3.3.6 software from Bruker AXS.

#### Synthesis of the iodine-labeled HA gel precursors

Firstly, HA-TIB **3a** and **3b** derivatives with a molar mass of 390 and 120 kg/mol, respectively, were synthesized by an amide coupling reaction between HA (0.200 g, 0.50 mmol) and *N*-(2-aminoethyl)-3-acetamido-2,4,6-triiodobenzamide (AcTIB-NH_2_, **2**) (0.177 g, 0.30 mmol) in a water/DMF (3/2, v/v) mixture containing DMTMM (0.10 g, 0.36 mmol) and the pH was adjusted to 6.5 using 1 M aqueous NaOH. The reaction was left for 48 h at room temperature under a vigorous stirring then, purified by ultrafiltration using deionized water. The iodine-labeled HA-TIB derivatives **3a** and **3b** were recovered by freeze-drying with 84 and 80 % yields, respectively. The DS of the HA-TIB derivatives **3a** and **3b** were found to be, respectively, 0.25 and 0.10 from ^1^H NMR analyses. In a second step, the derivatives **3a** and **3b** were reacted with fructosamine and APBA, respectively, according to the following conditions. For the synthesis of HA-TIB-Fru, fructosamine (0.012 g, 0.05 mmol) was added to a water/DMF (3/2, v/v) mixture containing DMTMM (0.090 g, 0.32 mmol) and HA-TIB **3a** (0.18 g, 0.32 mmol) and the pH was adjusted to 6.5 using 0.5 M aqueous NaOH. For the synthesis of HA-TIB-PBA, APBA (0.007 g, 0.036 mmol) was added to a water/DMF (3/2, v/v) mixture containing DMTMM (0.100 g, 0.36 mmol) and HA-TIB **3b** (0.16 g, 0.36 mmol) and the pH was adjusted to 6.5 using 0.5 M aqueous NaOH. After stirring for 24 h at room temperature, both HA derivatives were purified by ultrafiltration (membrane MWCO 10kDa) using deionized water and were recovered by freeze-drying with 90 % yield. The DS_Fru_ of the HA-TIB-Fru derivative **5** was found to be 0.15 and the DS_PBA_ of the HA-TIB-PBA derivative **7** was found to be 0.10 from ^1^H NMR analyses.

#### Gold nanoparticles synthesis

11-Mercaptoundecanoic acid-capped gold nanoparticles (11-MUDA AuNPs) were synthesized using a modified Turkevich method as previously described^[31]^. The procedure involved dissolving 85 mg of gold chloride salt in 500 ml of pure water, heating it while stirring until reaching boiling point, adding 25 mL of sodium citrate, boiling for another 15 minutes, and cooling to room temperature, resulting in a red wine-colored solution of gold nanoparticles. To cap the nanoparticles, 2.6 mg of 11-MUDA dissolved in 1 mL of ethanol was added and stirred overnight. The 11-MUDA AuNPs were then purified by centrifuging and exchanging the supernatant with pure water repeated 3 times. They were sterilized through syringe filtration with a size of 0.45 µm before use. These nanoparticles had a peak absorbance of 524 nm, average hydrodynamic diameter of 22 nm with a PDI of 0.2, core size of 11 ± 1 nm, and zeta potential of −44.4 mV.

#### Preparation of the HA-I and HA-ref hydrogels for rheometry

The HA-I hydrogel was prepared by mixing solutions of HA-TBA-PBA **7** and HA-TBA-Fru **5** in PBS (pH 7.4) at a total polymer concentration of 15 g/L (corresponding to a “pure HA” concentration of 12 g/L) and with a boronic acid/sugar molar ratio of 1/1, using a double-barrel syringe equipped with an extruder (MEDMIX, Switzerland). The hydrogel was directly transferred to plate of the rheometer. The HA-ref hydrogel was prepared using similar conditions, by mixing solutions of HA-PBA and HA-Fru in PBS (pH 7.4) at a total polymer concentration of 12 g/L (corresponding to a “pure HA” concentration of 12 g/L).

#### Agarose gel preparation and injection tests

Agarose gels were prepared by solubilizing agarose (300 mg) in 50 mL of PBS (pH 7.4) under stirring at 95 °C for 10 min. The agarose solution was then poured in an Eppendorf^®^ tube and the sample was kept at 4 °C for 24 h before the injection tests. The latter were carried out with a TJ-1A syringe pump controller (Aniphy, USA), at a rate of 5 µL/min).

#### Rheology

Dynamic rheological experiments were performed using a strain-controlled rheometer (ARES-RFS from TA Instruments) equipped with two parallel plates. All the dynamic rheological data were checked as a function of strain amplitude to ensure that the measurements were performed in the linear viscoelastic region. The parallel plate on which samples were placed has a diameter of 25 mm. The distance between the plates was 0.25 mm. A thin layer of low-viscosity silicone oil (50 mPa.s) was applied on the exposed surface of the samples, to prevent water evaporation. The details of the rheological measurements were as follows: 1) oscillatory frequency sweep (0.01-10 Hz) experiments were performed within the linear viscoelastic range (strain fixed at 10 %) to determine the frequency dependence of the storage (G’) and loss (G”) moduli; 2) oscillatory amplitude sweep experiments at 1 Hz were carried out to determine the linear-viscoelastic range of the hydrogel networks and the yield stress. They were immediately followed by time sweep experiments at 1 Hz and a strain of 10 % (linear viscoelastic region) to monitor the recovery of the rheological moduli; 3) alternate step strain sweep tests consisted in applying alternating strain deformations of 10 and 800% with a duration of 3 and 2 min, respectively, at a fixed frequency (1 Hz).

#### 3D cell viability study and cell proliferation

Human ASCs used in this study were isolated from human fat tissues after surgeries then purified against any diseases and viruses. All experiments were performed using hASCs at passage P2-P3. hASCs were cultured onto T175 flasks to reach 90-95 % confluency in a ⍺-MEM supplemented with 3% platelet lysatelysate and 1% heparin 5000 U/mL without antibiotics (penicillin/streptomycin). Cells were then trypsinized, pelleted and re-suspended into a growth media for cell counting.

#### Live/Dead assay

An aliquot of growth media containing hASCs were then taken to obtain a final density of encapsulated cells in the hydrogels of 5×10^5^ cells/mL. hASCs were pelleted and resuspended in a solution of the HA-TBA-Fru derivative in Dulbecco’s phosphate buffer 1X (DPBS, pH 7.4). The HA-TIB-PBA solution was transferred to Transwell^®^ cell culture inserts (0.4 µm porous membrane; Corning^®^, US) contained in 24-well plates. The cell-laden solution of HA-TIB-Fru in DPBS (pH 7.4) was pipetted and quickly transferred individually to the insert then mixed with the HA-TIB-PBA to form the cell-embedded iodine-labeled HA-I hydrogel. The cell-laden HA-ref hydrogel based on HA-PBA and HA-Fru with a final density of 5×10^5^ hASCs was prepared following the same procedure. The cell-laden hydrogels were supplemented with standard growth media and incubated for evaluating cell viability over 3 and 7 days at 37 °C – 5 % CO_2_. For each hydrogel condition, the experiment was carried out in triplicate (n=3) and with three different hASCs batches, at different times.

#### Live/Dead assay - image acquisition

A staining Live/Dead solution was made according to supplier information, by adding 10 μL of the Calcein-AM solution (green, excitation/emission 490/515 nm) and 5 μL of the propidium iodide solution (red, excitation/emission 535/617 nm) in 5 mL DPBS 1X. After 3 or 7 days of cells incubated in the hydrogels, the cell media was removed and the staining solution was then added to the gels with incubation for 15 minutes at 37 °C. Hydrogels containing hASCs were then analyzed under an epifluorescence microscope (ZEISS Axiovert 200M) equipped with a CoolSnap HQ^2^ camera using MetaMorph 3.5 imaging software. Images of cells encapsulated in 3D in the hydrogels were acquired using a Nikon 10X objective. 2D images were captured from 10 random fields of view for each sample. Fluorescent labeled cells in green (live cells) and red (dead cells) were counted using an image analysis software, ImageJ (National Institute of Mental Health, Bethesda, MD, USA; imagej.nih.gox/ij/).

#### Trypan blue assay

Individual cell-embedded hyaluronic acid hydrogels were treated by addition of Dotarem (aqueous solution of gadoteric acid, Gerbet^®^) diluted in DPBS 1X solution at 37 °C-5 % CO_2_ for 30 minutes to induce gel-sol transition and pelleted. hASCs were washed two times with a DPBS 1X solution, and then counted using a Trypan Blue solution 0.4%. A final cell density in the hydrogels was calculated compared to the initial cell density at day 0. Cell proliferation was estimated by the percentage of live and dead cells of each hydrogel at day 3 or day 7. For each hydrogel, triplicate (n=3) cell countings were individually performed at different times and with different hASC batches.

#### Gold nanoparticles (AuNPs) labeling of hASCs

Briefly, hASCs at passage P2-P3 were cultured onto a T175 flask in a volume of 30 mL of ⍺-MEM cell growth media for one week in physiological conditions 37°C – 5% CO_2_, in order to reach at 90-95 % confluence. The growth medium was replaced every 48h. At day 6, hASCs were incubated with gold nanoparticles at 0.1 mg/mL for 18 h based on a protocol that provided high AuNPs uptake while maintaining cell viability^[10]^. On day 7, gold-labeled hASCs (AuNPs-hASCs) were trypsinized, washed with a hot saline buffer then pelleted after swinging-bucket centrifugations (1000 rpm – 5 min).

#### 3D encapsulation of AuNPs-hASCs in HA-I hydrogel

As described previously, AuNPs-hASCs were collected as cell pellets after swinging-bucket centrifugations. AuNPs-hASCs were gently resuspended in 300 µL of HA-TIB-Fru solution prepared in DPBS 1X. The cell-laden solution was then pipetted, quickly transferred to an Eppendorf containing the HA-TIB-PBA solution and then mixed form the cell-embedded HA-I hydrogel. The cell-laden HA-I hydrogel was then transferred into a 1 mL syringe with attached 26G Needle.

##### Animal experiments

All experimental procedures involving animals and their care were carried out in accordance with the European regulations for animal use (EEC Council Directive 2010/63/EU, OJ L 276, Oct. 20, 2010). Data are reported according to ARRIVE guidelines (Animal Research: Reporting of In Vivo Experiments). An acclimation period of at least 7 days was observed before the start of the study. Because our *ex vivo* and *in vivo* studies aimed to provide a proof-of-concept of imaging feasibility (and not a treatment effect), we did not calculate a priori sample size, randomize animals in treatment groups nor blind allocation and analysis. All animals were included in the studies.

##### Brain study

The brain study was approved by the local review boards of Cermep imaging platform where treatments were administered and brains imaged *in vivo* with micro-CT and SPCCT (“Comité d’éthique pour l’Expérimentation Animale Neurosciences Lyon:” CELYNE; CREEA#042; APAFIS agreement #7457-2016110414498389) and of ESRF where the brains were imaged *ex vivo* with SKES-CT (ETHAX CREEA#011, APAFIS agreement #29284-2016110414498389). Four adult male Sprague-Dawley rats (age at reception: 6-7 weeks, body weight: 250-300 g) were purchased from Janvier (France). The animals were housed in a temperature- and humidity-controlled environment (21 ± 3 °C), with a 12-hour light-dark cycle. They had free access to food and water and their cages were enriched with transparent, red-colored tunnels and hazelnut wood sticks.

Rats were anesthetized by breathing 3-4% isoflurane (ISO-VET, Piramal Healthcare, Morpeth, UK) in air and maintained under 1.5% isoflurane in air using a face mask and mounted in a stereotaxic apparatus (D. Kopf Instruments). Isoflurane anesthesia was selected since it allows modulating anesthesia depth. Rectal temperature was kept at 37 ±1 °C throughout the surgical procedures, using a feedback-regulated heating pad. To alleviate pain, buprenorphine was administered subcutaneously at the dose of 0.05 mg/kg subcutaneously prior to the surgery. Lubricant ophthalmic gel was applied on both eyes to prevent ocular dehydration, and the surgical site was shaved and disinfected with antiseptic solution (i.e. 10 % iodopovidone). All rats had received one intracerebral injection in each of their two hemispheres (0.5 mm anterior-posterior (AP) to bregma, ±3.0 mm in medium-lateral (ML) direction, 5.5 mm dorsal-ventral (DV) from the cortical surface using a 10 µL Hamilton^®^ syringe with a 26G (0.4 mm) needle, beveled 40° (Hamilton, 701N, ThermoFischer, USA). The injection speed was calibrated at 2 µL/min with a waiting of 1 minute before the withdrawal of the 10-µL-precise syringe to prevent a backflow. Treatment consisted in 10µL vehicle (0.5×10^6^Au-hASCs in PBS solution, 10 µL, N=2 injections) or HA-I hydrogel (10-µL, N=2 injections) or the repair kit (0.5×10^6^ Au-hASCs in 10 µL HA-I hydrogel, N=4 injections). Treatment administration was followed by delivery monitoring using a micro-computed tomography (micro-CT). All rats were allowed to rest and to wake-up in their cage with an enriched food and water *ad libitum*. At day 3, heads were collected and prepared for an *ex vivo* bicolor X-ray monitoring with synchrotron k-edge subtraction computed tomography.

##### Knee study

The knee study was approved by the local review boards of the Languedoc-Roussillon Regional Ethics Committee on Animal Experimentation (approval CEEA-LR-10041, APAFIS agreement ##35861-2022031115332865) and GIN (CREEA#004”, APAFIS agreement #7457-2016110414498389) where the osteoarthritis model was performed and treatments were administered, and of ESRF where knees were imaged with SKES-CT *ex vivo* and *in vivo* (ETHAX CNREEA#011, APAFIS agreement #31781-20210520132410). Fifteen C57Bl/6J mice (age at reception: 1010 weeks, body weight: 20-25 g) were purchased from Charles River Laboratory (BerthevinL’Arbresle, France). The animals were housed in a temperature- and humidity-controlled environment (21 ± 3 °C), with a 12-hour light-dark cycle, free access to food and water and nest material according to the involved animal welfare units.

Collagenase-induced OA was induced as previously described^[32]^. In brief, right knee joints of mice were injected with 1 U type VII collagenase from *Clostridium histolyticum* (Sigma-Aldrich) in 5 μL of saline at day 0 and day 2, causing disruption of the ligaments and local instability of the joint. All surgery was performed under isoflurane gas anesthesia, and all efforts were made to minimize suffering. For the first experiments (*ex vivo* imaging), healthy mice (N=4) received intraarticular injections of AuNPs-hASCs embedded in HA-I hydrogel (2.5×10^5^ cells/ 2.5 µL of HA-I hydrogel: N=2 and 3.75×10^5^ cells/2.5 µL of HA-I hydrogel: N=2). Mice were sacrificed immediately post-injection and the joints were collected, fixed in ethanolic solution then secured in an agarose 1% hydrogel for *ex vivo* imaging. All injections were performed under isoflurane gas anesthesia. For the second experiment (*in vivo* imaging), a group of 11 mice with OA received AuNPs-hASCs embedded in HA-I hydrogel (2.5×10^5^ cells/ 2.5 μL HA-I hydrogel) at day 7 post-induction. Bicolor imaging with SKES-CT was performed on lived animals in the first 72h following administration (24h: N=2, 48h: N=4 and 72h: N=5). After the last imaging sessions, the joints were collected, fixed in 4% formaldehyde then embedded in 1% agarose gel for ex-vivo X-ray imaging with XPCT.

##### Multimodal computed tomography imaging

###### Micro-CT

Rats were imaged *in vivo* using a Preclinical microPET / CT camera (Siemens Inveon, 2010) on the Cermep imaging platform. The parameters used are 80kVp and 500 µA, 720 projections on 360° with an exposure time of 900 ms per projection, reaching an isotropic resolution of 55.6 µm.

###### SKES-CT

The SKES-CT acquisitions were performed on the biomedical beamline ID17 of the European synchrotron radiation facility (ESRF). The high flux of photons from synchrotron sources allows to perform monochromatic acquisitions using a double bent Laue monochromator (ΔE/E = 0.1%). Samples and animals were imaged at multiple energies around the K-edge of materials of interest present in the imaged subject. When iodine is present in the sample, two acquisitions are made around the K-edge of iodine (33.17 keV) at 32.2 keV and 34.2 keV. In case of presence of gold, two acquisitions are made around the K-edge of gold (80.7 keV) at 79.7 keV and 81.7 keV. The position of the samples being far from the detector (3.5 m) and the beam having a strong spatial coherence, it was possible to use propagation-based phase contrast to enhance contrast in the images. The *ex vivo* knee samples were imaged with an isotropic resolution of 6.5 µm. Rat brains as well as phantoms were imaged with an isotropic resolution of 21.3 µm. Mice imaged *in vivo* reached an isotropic resolution of 13.3 µm.

##### Material decomposition

Material decomposition process allows to obtain concentration maps of the elements of interest (gold and iodine), by using monochromatic images of the same sample at different energies. Using two images and making the hypothesis that each voxel is composed of three elementary materials M_i_ (M_G_: gold, M_I_: iodine and M_W_: water), we just have to resolve the following system:

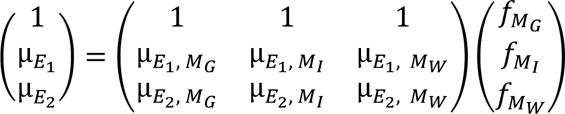

With *E*: the energy of the X-ray beam [keV]

µ_E,_*M_i_*: the linear attenuation coefficient of the material *i* at the energy E [cm^−1^]

f_M_i__: the volume fraction occupied by the element *i* within a voxel

This system admits a unique solution 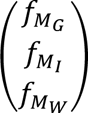, corresponding to the concentrations of the materials of interest in each pixel of the sample image. The accuracy of such measurement greatly depends on the materials concentration and the X-ray dose.

##### XPCT

XPCT acquisitions were done with a “pink” incident X-ray beam generated by the undulator on ID19 beamline (energy: 26 keV). The Paganin single distance phase-retrieval approach was used for data reconstruction reaching an isotropic resolution of 6.5 µm.

##### Segmentation method

The segmentation method is described in detail in a previous work^[33]^. Briefly, a thresholding technique is used. The gold threshold is 0.75 mg/mL and the iodine one is 0.25 mg/mL. Morphological opening with structuring element of radius 2 pixels is performed. For the brain study we keep only the segmented within the cranial skull and for the knees we performed a connected component analysis to keep only relevant objects.

## Acknowledgments

This project was funded by the French national research agency (ANR18-CE19-003, Breakthru grant), and partly funded by the GlycoAlps IDEX UGA in the framework of the Investissements d’avenir program [ANR-15-IDEX-02]). ANR also supported the national infrastructure: “ECELLFRANCE: Development of a national adult mesenchymal stem cell based therapy platform” (ANR-11-INSB-005). We thank Caroline Bouillot and Salim Si-Mohamed for performing respectively µCT and SPCCT experiments at Lyon’s multimodal imaging platform Cermep. The authors thank the NMR platform of ICMG (FR2607) for its support; the European Synchrotron Radiation Facility (ESRF, beamline ID17) for allocation of beamtime (MD1333) and their local contact Herwig Requardt for help during the experiments. DPC acknowledges the NIH for support (R21-EB029158 and R21-EB029556).

## Data Availability Statement

Imaging data are available under reasonable request addressed to the corresponding author.

## Authors’ contributions (CRediT)

**Conceptualization:** EBa, CJ, DN, DC, EBr, HE, MW, OD, CR, RA

**Data Curation:** MS, CT

**Formal analysis:** MS, CT

**Funding acquisition:** DC, EB, MW

**Investigation:** MS, CT, CD, KT, YCD, NC, CA, AM, BF, BC, DN, EB, HE, MW, CR, RA

**Methodology:** EBa, CJ, DN, DC, EBr, HE, MW, OD, CR, RA

**Project administration:** EB, HE, MW, OD, CR, RA

**Resources:** YCD, BC, EBa, CJ, DC, CA, AM

**Software:** EB, CT

**Supervision:** DN, DC, EB, HE, MW, OD, CR, RA

**Validation:** EB, MW, RA

**Visualisation:** MS, CT, EB

**Writing, original draft:** MS, CT, MW, RA

**Writing, review and editing:** All authors

