## Supporting Information for "A novel injectable radiopaque hydrogel with potent properties for multicolor CT imaging in the context of brain and cartilage regenerative therapy"

(d) Univ. Grenoble Alpes, Inserm, UA7 Strobe, 38000 Grenoble, France

(e) Hospices Civils de Lyon, France

(f) IRMB, Univ. Montpellier, INSERM, CHU Montpellier, Montpellier, France

(g) Department of Radiology and Department of Bioengineering, University of Pennsylvania, Philadelphia, Pennsylvania 19104, United States

(h) Hôpital Edouard Herriot, Lyon, France

(i) Cell Therapy and Engineering Unit, EFS Rhône Alpes, 38330 Saint Ismier, France

(j) Univ. Grenoble Alpes, Translational Innovation in Medicine & Complexity, UMR552, 38700 La Tronche, France

(k) Univ. Grenoble-Alpes, Département de Pharmacochimie Moléculaire UMR 5063, 38400 Grenoble, France

Institut de Biologie et Pathologie, CHU de Grenoble-Alpes, 38700 La Tronche, France

(l) CHU Grenoble Alpes, Stroke Unit, Department of Neurology, 38043 Grenoble, France

†Equal contribution

**Table of Contents**

1. Synthesis of *N*-(2-aminoethyl)-3-acetamido-2,4,6-triiodobenzamide (AcTIB-NH_2_)

2. ^1^H NMR spectra of AcTIB-NHBoc, AcTIB-NH_2_, HA-TIB-Fru and HA-TIB-PBA derivatives

3. Rheological analysis of the HA-ref hydrogel in PBS at pH 7.4

4. Cell viability of human adipose-derived stem cells in 2D cell

5. Cytotoxicity (MTT) assay of individual solutions of HA derivatives incubated with hADSCs

References

**1. Synthesis of *N*-(2-aminoethyl)-3-acetamido-2,4,6-triiodobenzamide (AcTIB-NH_2_)**

3-Acetamido-2,4,6-triodobenzoic acid bis(2-hydroxyethyl)-ammonium salt, (0.2 g, 0.3 mmol) and HATU (0.229 g, 0.60 mmol) were dissolved at room temperature in 50 mL of anhydrous *N,N*-dimethylformamide. After stirring for 10 min under nitrogen, *N*-Boc-ethylenediamine (0.096 g, 0.6 mmol) and diisopropylethylamine (0.234 g, 0.0 mmol) were added and the reaction medium was stirred overnight at room temperature under nitrogen. After evaporation of the solvent under reduced pressure, the crude product was dissolved by addition of ethyl acetate. The organic phase was washed thoroughly with 0.5 M HCl, aqueous saturated NaHCO_3_, and brine, dried over anhydrous sodium sulphate, filtered through filter paper and finally, concentrated by rotary evaporation. *N*-Boc-(2-aminoethyl)-3-acetamido-2,4,6-triiodobenzamide **(AcTIB-NHBoc, Figure S1)** was obtained as a white powder in 85 % yield. The structural integrity and purity of the crude product was confirmed by ^1^H NMR spectroscopy (**Figure S2**).

AcTIB-NHBoc (0.180 g, 0.26 mmol) was then dissolved in a mixture dichloromethane/methanol (1 mL, 9/1, v/v) and excess TFA (5 mL, 65 mmol) was added. After stirring for 4 h at room temperature, the solvent and TFA are removed by rotary evaporation to yield AcTIB-NH_2_.

^1^H NMR (400 MHz, DMSO-d6+D_2_O, 25 °C) of AcTIB-NH_2_: δ_H_ (ppm) 9.94 (1H, H_Ar_), 3.44 (2H, C*H_2_*NHCO), 2.99 (C*H_2_*-NH2), 2.02 (3H, CH_3_CO).

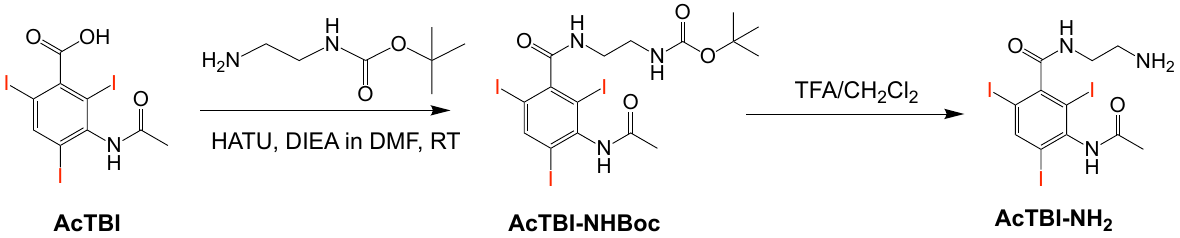

**Figure S1.** Reaction conditions for the synthesis of AcTIB-NH_2_.

**2. ^1^H NMR spectra of AcTIB-NHBoc, AcTIB-NH_2_, HA-TIB-Fru and HA-TIB-PBA derivatives.**

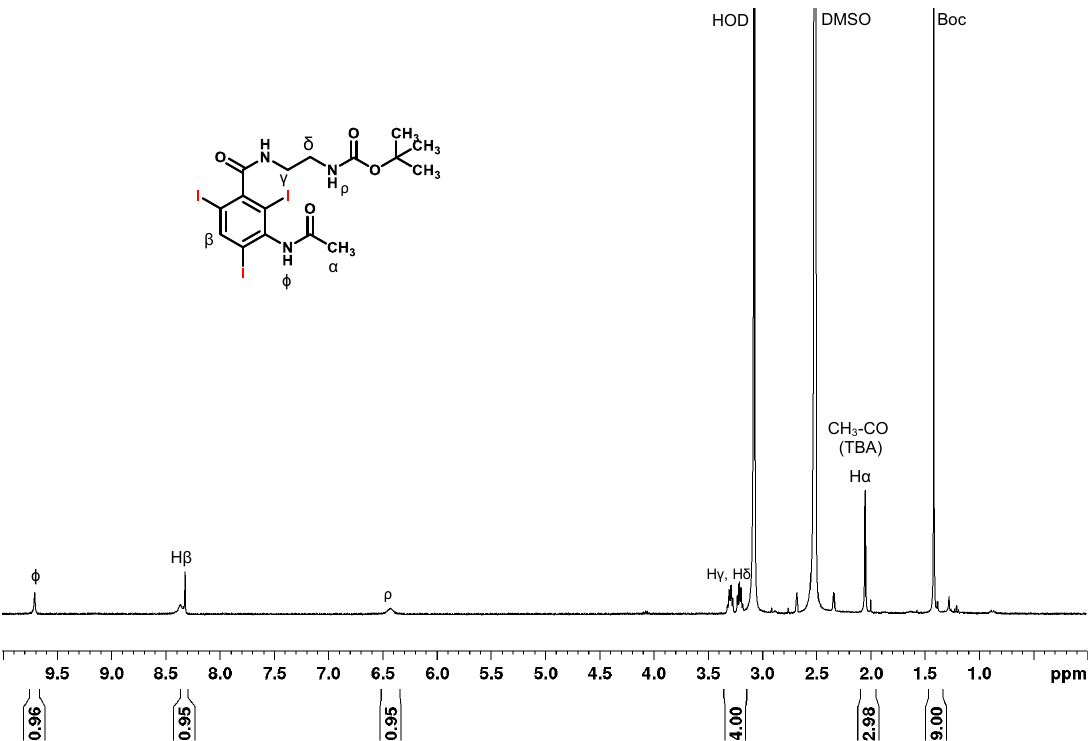

**Figure S2.** ^1^H NMR spectrum (400 MHz, DMSO-d6, 6 mg/mL, 25 °C) of *N*-Boc-(2-aminoethyl)-3-acetamido-2,4,6-triiodobenzamide (AcTIB-NHBoc).

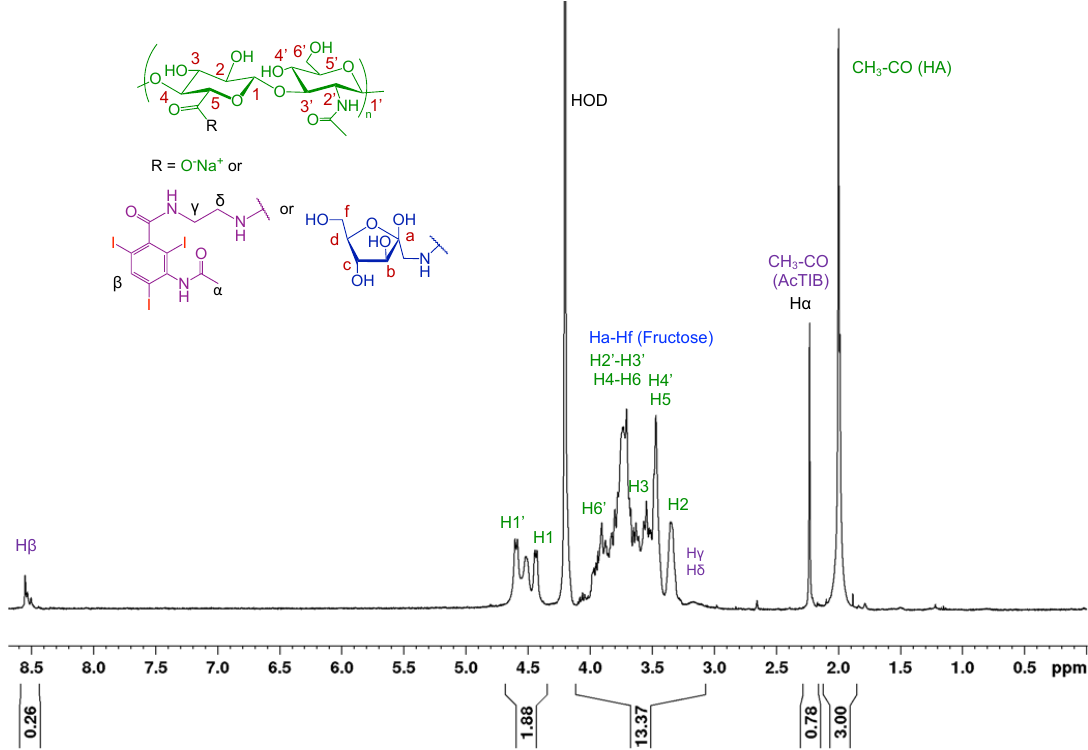

**Figure S3.** ^1^H NMR spectrum (400 MHz, D_2_O, 6 mg/mL, 80 °C) of HA-TIB-Fru.

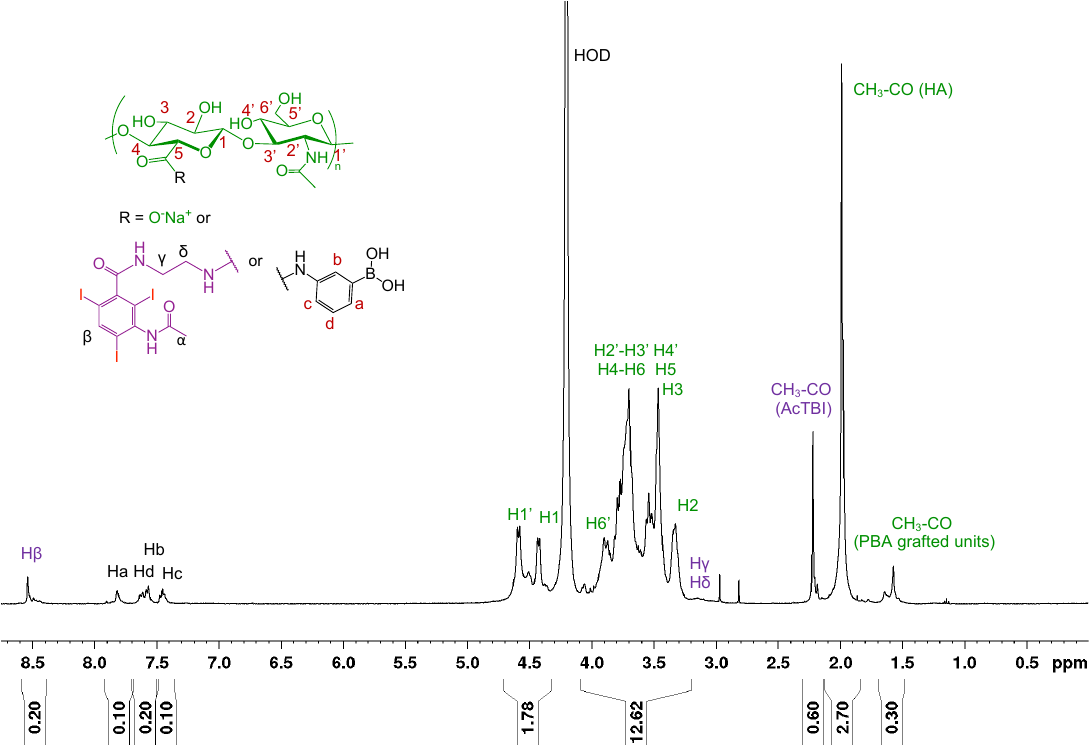

**Figure S4.** ^1^H NMR spectrum (400 MHz, D_2_O, 6 mg/mL, 80 °C) of HA-TIB-PBA.

**3. Rheological behavior of the HA-ref hydrogel in PBS, pH 7.4**

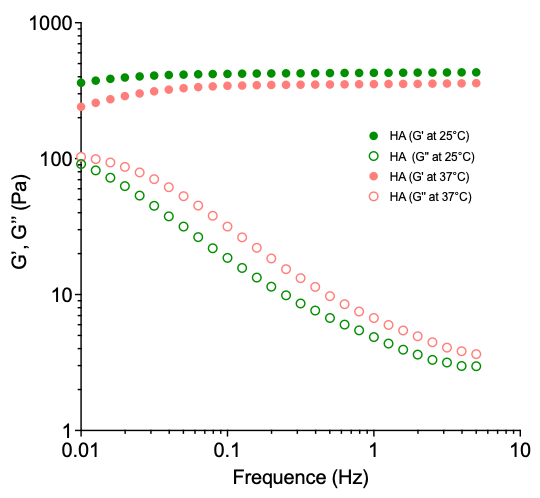

**Figure S5.** Dynamic rheological behavior of the HA-ref hydrogel (PBA/Fru molar ratio of 1) prepared from HA-PBA and HA-Fru in PBS, pH 7.4 (*C_p_* = 12 g/L). Frequency dependence of the storage modulus (G’, filled symbols) and loss modulus (G’’, empty symbols) measured with 5% strain at 25°C and 37°C.

**4. Cell viability of human adipose-derived stem cells in 2D cell culture**

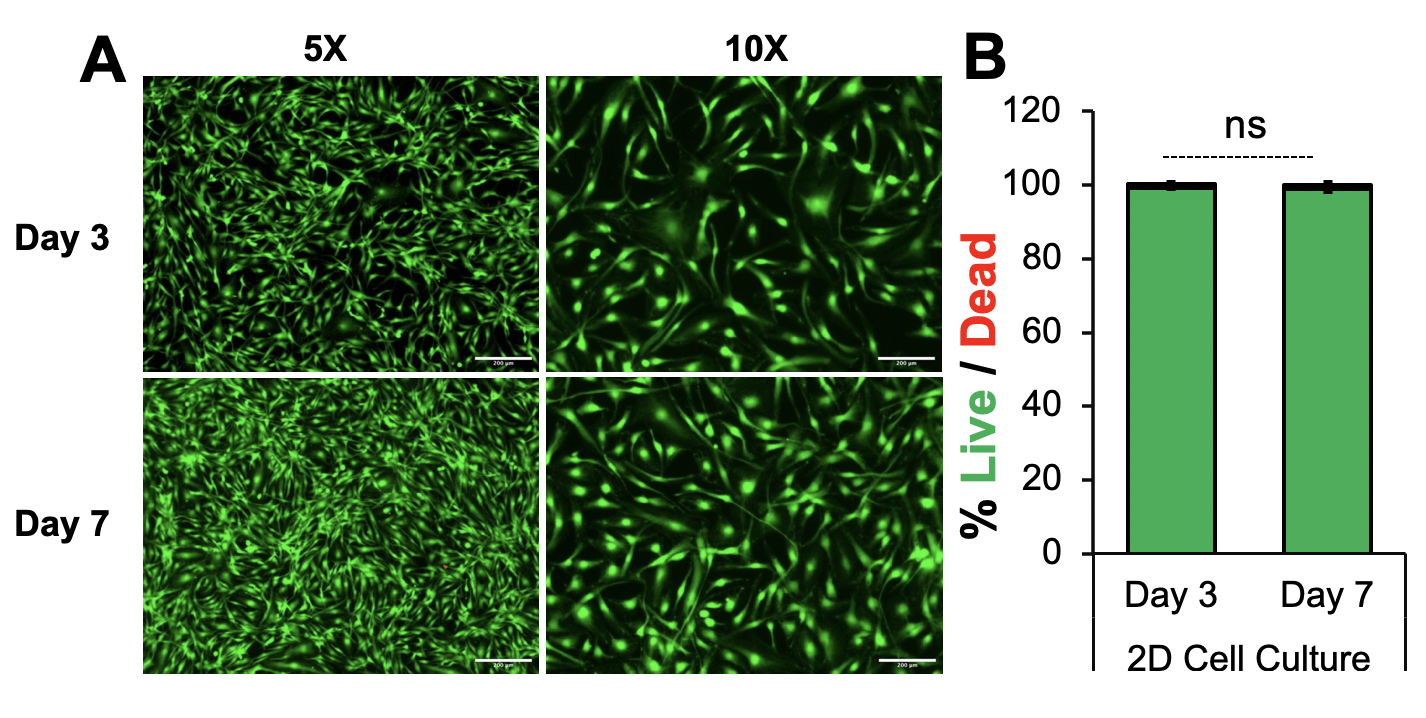

**Figure S6.** Cell viability of hASCs harvested in 2D cell culture at day 3 and day 7 in a MEM-α cell culture medium supplemented with 1% lysat plaquet and 1% heparin without antibiotics (penicillin/streptomycin). Cells were stained with calcein AM (green, live cells) and ethidium homodimer-1 (red, dead cells) using a diluted Live/Dead kit solution prepared in DPSBS 1X (Sigma, France). **A**) Microscopy images of hASCs for the Live/Dead assay at day 3 and day 7, in taken with a ZEISS Axiovert 200M epifluorescent inverted microscope in a 5X and 10X objectives with a CCD CoolSnap HQ camera using MetaMorph 3.5. B) Cell viability of hASCs represented as mean ± standard deviation based on 3 different hASCs patient donors, cultivated on triplicate conditions at different cell culture periods. Statistical analysis was performed by one-way analysis of variance (ANOVA) with a Tukey post-hoc multiple comparisons tests with a 95% confidence interval (⍺ = 0.05, * p < 0.5, ** p < 0.01, *** p < 0.001, **** p < 0.0001, ns for non-significative value) using GraphPad 9.3.0 (San Diego, CA, USA).
